# Molecular mechanisms controlling the multistage post-translational processing of endogenous Nrf1α/TCF11 proteins to yield distinct proteoforms within the coupled positive and negative feedback circuits

**DOI:** 10.1101/300327

**Authors:** Yuancai Xiang, Meng Wang, Shaofan Hu, Lu Qiu, Fang Yang, Zhengwen Zhang, Siwang Yu, Jingbo Pi, Yiguo Zhang

## Abstract

In an attempt to terminate the chaotic state of the literature on Nrf1/TCF11 with various confused molecular masses, we herein establish a generally acceptable criterion required for identification of its endogenous full-length proteins and derivative isoforms expressed differentially in distinct experimental cell lines. Further work has been focused on the molecular mechanisms that dictate the successive multistate post-translational modifications (i.e. glycosylation by OST, deglycosylation by NGLY, and ubiquitination by Hrd1) of this CNC-bZIP protein and its proteolytic processing to yield multiple isoforms. Several lines of experimental evidence have demonstrated that the nascent Nrf1α/TCF11 polypeptide (non-glycosylated) is transiently translocated into the endoplasmic reticulum (ER), in which it becomes an inactive glycoprotein-A, and also folded in a proper topology within and around membranes. Thereafter, dynamic repositioning of the ER-resident domains in Nrf1 glycoprotein is driven by p97-fueled retrotranslocation into extra-ER compartments. Therein, glycoprotein of Nrf1 is allowed for digestion into a deglycoprotein-B and then its progressive proteolytic processing by cytosolic DDI-1/2 and proteasomes to yield distinct proteoforms (i.e. protein-C/D). The processing is accompanied by removal of a major N-terminal ^~^12.5-kDa polypeptide from Nrf1α. Interestingly, our present study has further unraveled that coupled positive and negative feedback circuits exist between Nrf1 and its cognate target genes, including those encoding its regulators p97, Hrd1, DDI-1 and proteasomes. These key players are differentially or even oppositely involved in diverse cellular signalling responses to distinct extents of ER-derived proteotoxic and oxidative stresses induced by different concentrations of proteasomal inhibitors.

## INTRODUCTION

The transcription of genetic information from genomic DNAs to messenger RNAs is controlled by a group of proteins (i.e. transcription factors) through binding specific *cis*-regulatory sequences in their target gene promoters in order to monitor a programme of increased or decreased gene expression [1, 2]. As such transcription factors are active, they play versatile roles essential for many cellular processes, including cell cycle, division, differentiation, metabolism and defence [3–5]. Notably, a CNC-bZIP (cap‘n’collar basic-region leucine zipper) family of transcription factors comprises the *Drosophila* CNC protein [6], the *Caenorhabditis elegans* Skn-1 [7], the vertebrate NF-E2 (nuclear factor-erythroid 2) p45 subunit and related factors Nrf1 (with its long TCF11 and short Nrf1β/LCR-F1 forms), Nrf2 and Nrf3 [8, 9], in addition to Bach1 and Bach2 [10]. Except Skn-1, all other CNC-bZIP proteins heterodimerize with small Maf or other bZIP proteins prior to binding to antioxidant response elements (AREs) or homologous sequences in their target genes. ARE-driven genes regulated by CNC-bZIP factors include those encoding antioxidant proteins, detoxification enzymes, metabolic enzymes and 26S proteosomal subunits [8, 9, 11], which are responsible for critical homeostatic, developmental and cytoprotective pathways. In the response to cellular stress, basal and inducible transcriptional expression of these ARE-driven genes is finely-tuned, in order to maintain cellular homeostasis and organ integrity. In fact, it is true that the adaptive defence response enables transactivation of other ARE-battery genes, such as those encoding DNA repair enzymes, cofactor-generating enzymes, molecular chaperones and anti-inflammatory response proteins [12–14]. However, a chronic long-term redox stress could over-stimulate expression of key genes involved in regulation of cell-cycle, metabolism, immune response, neurodegeneration, apoptosis, necrosis, inflammation and other pathobiological processes. Such pathological over-stimulants clearly contribute to carcinogenesis [15, 16] and ageing-related neurodegenerative [17–19].

In mammals, Nrf1 and Nrf2 are two principal CNC-bZIP factors to regulate ARE/EpRE-driven cytoprotective genes against cellular stress [20, 21]. Accumulating evidence has revealed that the membrane-bound Nrf1 fulfils a unique biological function, that is distinctive from the water-soluble Nrf2, in maintaining cellular homeostasis and organ integrity, although Nrf2 is widely accepted to be a master regulator of adaptive responses to oxidative stressors and electrophiles [22, 23]. The latter notion is drawn from the fact that global *Nrf2* knockout (*Nrf2*^−/−^) mice are more susceptible than wild-type animals to carcinogens and oxidative stress [24], but they have not developed any spontaneous cancer or diabetes [25]. By contrast, gene-targeting experiments of *Nrf1* (called *nfe2l1*) demonstrate that it, unlike Nrf2, is indispensable for normal development and healthy growth, because global knockout of *Nrf1* in the mouse causes embryonic lethality [26, 27] and severe oxidative stress [28, 29]. Further, conditional knockout of *Nrf1* in specific organs results in distinct pathological phenotypes, such as non-alcoholic steatohepatitis (NASH)-based hepatoma [30, 31] and neurodegenerative diseases [32, 33]. Therefore, Nrf1 is taking great strides into its own *bona fide* significances in order to become a hot and attractive theme. However, the underlying mechanisms by which the unique functioning of Nrf1 is tightly regulated at various levels have not been well elucidated [9, 34].

To date, ever-increasing evidence has revealed that the functional activity of Nrf1 is finely tuned by a steady-state balance between its production and the concomitant processing into distinct isoforms (called proteoforms) before being turnover [11, 35–37]. These proteoforms are together coordinated to confer on the host robust cytoprotection against a variety of cellular stresses. In distinct topovectorial processes after translation of ectopic Nrf1 factor [38, 39] (Figure 1A), it is allowed for the selective post-synthetic processing of the CNC-bZIP protein in different tempo-spatial subcellular locations from the endoplasmic reticulum (ER) to the nucleus [21, 40–44]. Consequently, Nrf1 is modified successively by N-glycosylation, O-GlcNAcylation, deglycosylation, phosphorylation, ubiquitination, degradation and proteolytic cleavage. This leads to yield of multiple distinct isoforms (i.e. 120-kDa, 95-kDa, 85-kDa, 55-kDa, 46-kDa, 36-kDa and 25-kDa estimated by NuPAGE), which exert different and even opposing transcriptional activities to mediate differential target gene expression [45–48]. Amongst isoforms, some are also generated from alternative translation starting signals located within various lengths of multiple mRNA transcripts (to yield TCF11, Nrf1α, Nrf1β and Nrf1γ) [49–51]. The alternative splicing of some transcripts, arising from distinct transcription start sites existing within the single *Nrf1* gene loci, leads to the translation of several deletion mutants (i.e. Nrf1D, Nrf1^ΔN^ and Nrf1^ΔS^) [52, 53]. Nonetheless, much remains unknown about the detailed mechanisms underlying alternative translation and transcription of Nrf1 to yield multiple endogenous products from its single *Nfe2l1* gene (with 23 transcripts including splicing variants, along with 19 encodable proteins, deposited in the Ensembl Genome).

**Figure 1.**
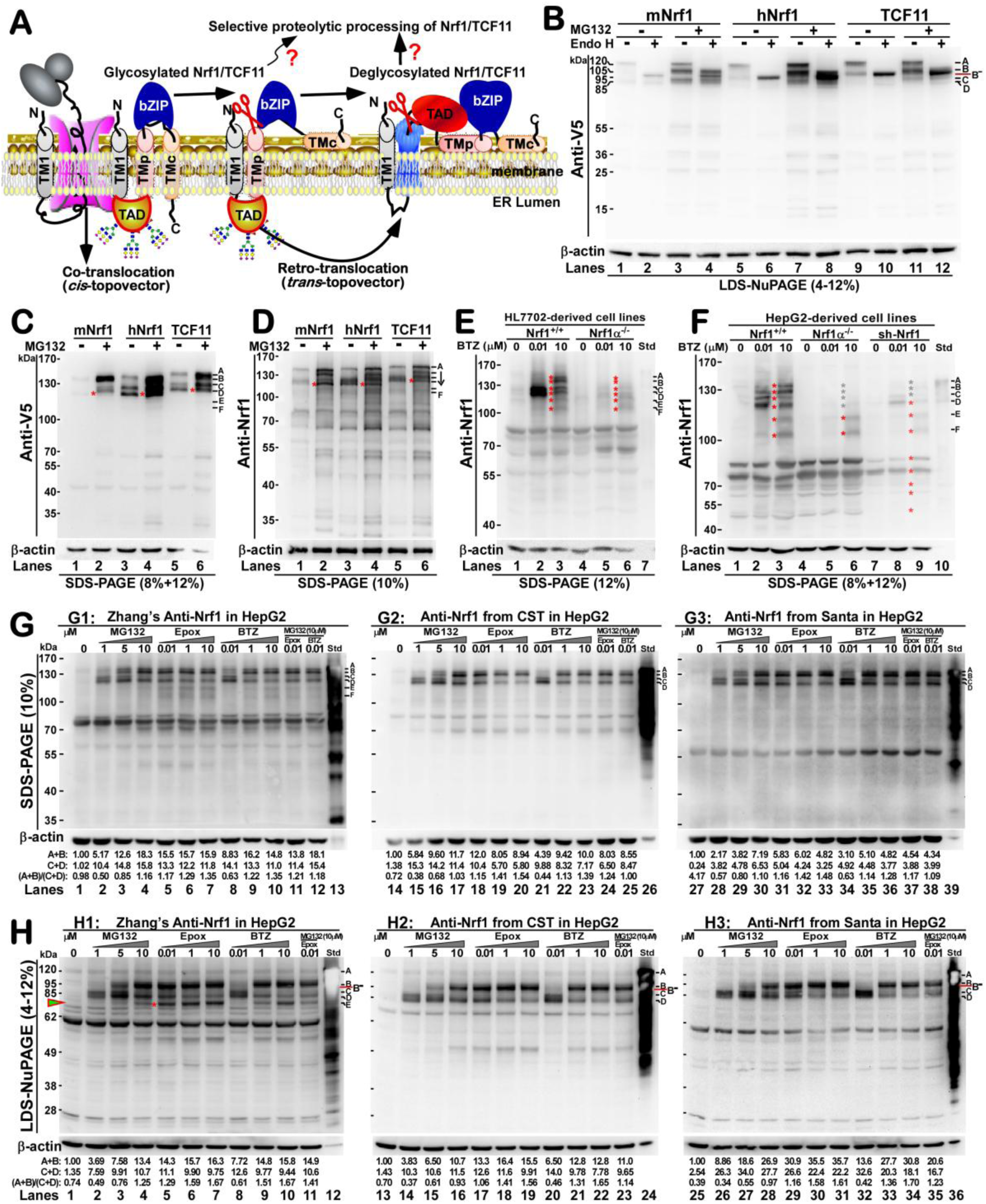
Multi-strategic identification of electrophoretic locations of distinct Nrf1 isoforms. **(A)** Schematic representation of the biosynthesis and putative post-translational processing of Nrf1, that is entailed by dynamic topologies across ER membranes. **(B to D)** Expression constructs for mouse (mNrf1) and human (hNrf1) full-length proteins along with human TCF11 (all three were tagged C-terminally by the V5 ectope) were transfected into COS-1 cells for 7 h before being allowed for a 16 h recovery. Subsequently, the cells were treated with 5 μmol/L of MG132 for 4 h (B) or 16 h (C, D) before being harvested in a denature buffer. Then total lysates were allowed for deglycosylation reactions with Endo H for 4 h at 37°C, prior to being resolved by 4-12% NuPAGE gels and visualized with V5 antibody (B). Similar samples, without incubation of Endo H, were subjected to protein separation by SDS-PAGE gels containing distinct concentrations of polyacrylamide, followed by immunoblotting with antibodies against V5 (C) or Nrf1 (D), respectively. **(E,F)** Both HL7702 and HepG2 (*Nrf1^+/+^*), as well as their derived Nrf1α-specific knockout (Nrf1α^−/−^) and stable sh-Nrf1-expressing, cell lines were treated with the indicated doses (0-10 μmol/L) of BTZ for 16 h before being harvested in denature buffer. The total lysates were then isolated by SDS-PAGE containing different concentrations of polyacrylamide and visualized by Western blotting with an Nrf1-specific antibody (saved in our lab). In addition, a positive standard (Std) sample was referred to lysates of cells that had been transfected with an expression construct for full-length Nrf1. **(G,H)** HepG2 cells were treated for 16 h with different doses of proteasomal inhibitors MG132, Epox and BTZ. Equal protein amounts of these total lysates were loaded in each well of SDS-PAGE gels containing 10% polyacrylamide (G) or gradient LDS-NuPAGE gels containing 4-12% polyacrylamide (H), before being transferred to the PVDF membrane and then visualized by immunoblotting with distinct antibodies against Nrf1 (i.e. Zhang’s lab and other two antibodies from Cell Signaling Technology and Santa Cruz Biotechnology). The relative intensity of major protein bands and their ratios were quantified by using the Quantity-one software and shown on the bottoms.

To gain an in-depth insight into the molecular basis for different *Nrf1*^−/−^ phenotypes (in which certain Nrf1 species were deleted or retained), a great task is to identify differential abundances of distinct molecular weight isoforms of endogenous proteins existing in model cell lines. Unfortunately, most of different experimental settings have given rise to controversial and even self-contradictory results published in the current literature. Firstly, only p60 protein was not detected (but polypeptides of ^~^95, 65, 50 and 48-kDa were retained) by western blotting of the *Nrf1*^−/−^ lyastes with the primary peptide antibody against aa 565-586 of Nrf1 (i.e. aa 274-293 in Nrf1β/LCR-F]/p65Nrf1) [27]. Secondly, another rabbit antibody against Nrf1 (purified from ant-Nrf1/GST fusion sera and identified by [41] in a cropped Figure S1F) was employed for western blotting of the same *Nrf1*^−/−^ lyastes as described above, showing that both p120 and p65 proteins were undetectable, with an exception of both minor p110 and major p60 proteins being retained [54]. Further examinations of ARE-battery genes as *Nqo1*, *Gclc*, *Gsta2*, but not *Gclm*, were up-regulated mouse embryonic fibroblasts (MEFs) derived from *Nrf1*^−/−^ animals, though they lacked p120 and p65 [54]; this appears to contradict their previous claims [29]. Thirdly, the rabbit polyclonal antibody against the N-terminus of Nrf1 (raised in the Chan laboratory [27, 54]) was subjected again to Western blotting, revealing that there still expressed a major extra ^~^85-kDa protein and two minor isoforms of p120 and p110, together with another two proteins closely to p65, but not the exact p65, being retained to considerably lowered levels in *Nrf1*^−/−^ MEFs [35]. All these remaining proteins were further abolished by the retrovirus expressing sh-Nrf1 (short hairpin-RNA interfering Nrf1), despite no changes in a small polypeptide recognized by a C-terminus-specific antibody (Novus Biologicals), apart from the fact that inducible transcription of the *proteasome* (*PSM*) genes was abrogated by *Nrf1*^−/−^, but not *Nrf2*^−/−^ [35]. Fourthly, *Nrf1*^−/−^ (from the knock-in mutant *Nrf1*^*rPGK-neo*^) cells also enabled to express certain Nrf1 isoforms to varying extents, and such these resulting experimental data should be interpreted with extreme cautions. Such distinct Nrf1 isoforms are inferable to be putative by-products from a fraction of mRNA transcripts being alternatively spliced to remove certain portions of the knocked-in exogene-coding sequence or *en bloc*; their residual expression levels may lead to a disparity in the obvious pathophysiological phenotypes occurring in between different strategic knockout mice (cf *Nrf1*^*rPGK-neo*^ [27] with *Lcrf1^tm1uab^* [26]).

Such a hot subject of debate, with sensible doubtful points reported by the authors [37, 55–57], has been focused primarily on whether and how Nrf1/TCF11 is allowed for selective proteolytic processing of the CNC-bZIP proteins by putative Hrd1/p97-driven ERAD-dependent ubiquintin proteasome pathway [11, 36, 37, 46] or aspartic proteases DDI-1/2 [58, 59]. In the meantime, this debate has reflected a fact that mechanisms underlying dual opposing effects of proteasomal inhibitors on Nrf1-mediated transcription of proteasomal subunit genes remains elusive. In several studies, proteasomal dysfunction by its inhibitors was considered to result in inducible activation of Nrf1 (and Skn-1) by proteasome-limited proteolysis of the CNC-bZIP factor, particularly in the ‘bounce-back’ response to relative lower doses of proteasomal inhibitors [11, 35, 37, 60]. However, no available evidence has been provided showing that a sufficient blockage of the proteasome activity leads to a significant attenuation or complete abolishment in the putative proteolytic processing of Nrf1/TCF11 [11, 47, 56, 61, 62]. Intact glycoprotein and deglycoprotein of Nrf1 are estimated to be between 140-kDa and 120-kDa on the routine SDS-PAGE gels, but two similar isoforms are resolved by LDS-NuPAGE gels to migrate between 120-kDa and 95-kDa [39, 46]). This is at least worth arguing against a recent report by Sha and Goldberg [37], who assumed that the proteasome-limited protealytic processing of Nrf1/TCF11 protein with a mass of ^~^100-kDa to yield a cleaved ^~^75-kDa form, which was thought to be abolished by high doses of proteasomal inhibitors (Figure S1D). This resulted for protein separation by SDS-PAGE gels and immunoblotting with two distinct antibodies H285 and C19, that recognize the central peptide (aa 191-475) of human Nrf1/TCF11 and its C-terminus, respectively (from Santa Cruz Biotechnology Inc). Later, the cleaved Nrf1 isoform of ^~^75-kDa was also corrected to be approximately 90-95 kDa [55], which were recognized by D5B10, a monoclonal antibody against human Nrf1/TCF11 at its N-terminal G129-surrounding peptide that is not overlapped with the adjacent aa 191-475 peptide (from Cell Signaling Technology Inc, Figure S1E). Collectively, both whether and how human endogenous Nrf1/TCF11 proteins are post-translationally processed into multiple isoforms are well not understood.

To provide a clear better understanding of molecular mechanisms underlying the post-translational processing of endogenous Nrf1α/TCF11 to yield multiple proteoforms, we have determined: (i) whether its full-length protein and major derivatives between 140-kDa and 70-kDa are differentially expressed in various cell types, aiming to establish a general criterion acceptable for identification of distinct isoforms; (ii) which are generated from post-translational modifications by glycosylation, deglycosylation and ubiquitination; (iii) which proteoforms (with distinct half-lives and activities) arise from the Hrd1/p97-driven proteasome-mediated proteolysis or aspartic proteases DDI-1/2-mediated cleavage of the intact CNC-bZIP protein and its longer isoforms, upon treatments with different doses of proteasomal inhibitors; (iv) mechanisms underlying the dual opposing effects of proteasomal inhibitors on differential expression of Nrf1, that is activated in the intracellular ‘bounce-back’ response to lower doses of proteasomal inhibitors but also repressed by higher doses of the same chemicals; (v) whether positive or negative feedback circuits exist between Nrf1 and its target genes, that are involved in the multistage post-translational processing of the CNC-bZIP factor, critical for discrete cellular signalling responses to distinct concentrations of proteasomal inhibitors, which stimulate distinct extents of both ER-derived proteotoxic and oxidative stresses.

The above-described samples were also subjected to further protein separation by two parallel SDS-PAGE gels, that contain different concentrations of polyacrymide in the routine Tris-glycine SDS running buffer (pH 8.3), which is distinctive from those of LDS-NuPAGE in the MES SDS running buffer (pH 7.3) [46]. Subsequently, Western blotting of MG132-treated cell lysates with antibodies against the V5 tag (attached to the C-terminal end of Nrf1/TCF11) showed that ectopic mNrf1 proteins were represented by two major closer bands of about ^~^140-kDa (i.e. defined as A and B) and another tow minor bands of approximately ^~^120-kDa (i.e. defined as C and D) (Figure 1C, *lane 2*). Relatively, such four protein bands from A to D appeared to be clearly visualized by Nrf1-specific antibodies against its short isoform Nrf1β (aa 292-741, raised in Zhang’s own laboratory), of which these two bands B & D seemed to emerge following MG132 treatment (Figure 1D, *lanes 2 vs 1*). Such a subtle nuance in the recognition of the protein-D form by distinct antibodies against Nrf1β or its C-terminal V5 tag also suggests a possible difference in the C-terminal processing of mNrf1. By contrast, either hNrf1 or its long isoform TCF11 in untreated COS-1 cells was resolved to exhibit similar three major protein bands A, C and D, but treatment of cells with MG132 caused a clear electrophoretic pattern of their protein bands A, B, C and D (Figure 1, C & D). This demonstrates that the B-form is a transient unstable protein, and thus it is inferable that it may be proteolytically processed (and rapidly degraded) to disappear or to be replaced by other processed isoforms, particularly in the absence of the proteasomal inhibitors.

To identify which bands represent the endogenous proteins of human Nrf1α/TCF11 and their derivative isoforms, TALENs-mediated editing of its gene was employed to knock out a site-specific DNA sequence surrounding the first translation start codon for the full-length Nrf1α/TCF11 protein (i.e. *Nrf1α*^−/−^) in two cell lines of human hepatocellular carcinoma (HepG2, with almost none of TCF11 transcripts confirmed by sequencing) and the immortalized human liver HL-7702 (with an 1:1 ratio of Nrf1α to TCF11 transcripts detected). As anticipated, the above-aforementioned four major bands A to D (representing endogenous Nrf1α/TCF11 and their derivatives) were significantly prevented (Figure 1E) and even completely abolished (Figure 1F) in two different *Nrf1α*^−/−^ cell lines, even upon treatment of the cells with bortezomib (BTZ, another proteasomal inhibitor, that is distinctive from MG132). However, additional two bands between 115-kDa and 105-kDa (i.e. E & F -forms), as well as small isoforms of < 100-kDa, were unaffected by the deficiency of *Nrf1α*^−/−^, suggesting that they are not derivatives directly from the putative proteolytic processing of intact Nrf1α/TCF11. By coincidence, the parallel experiments revealed that all those proteoforms between 140-kDa and 45-kDa were significantly diminished by knockdown of sh-Nrf1 targeting to all their C-terminal region-encoding transcripts (Figure 1F, *lanes 7-9*). In addition, the endogenous mNrf1 isoforms of ^~^140-, 105-, 68-, and 55-kDa were also markedly blocked by knockout of *Nrf1*^−/−^ in MEFs (Figure S2A).

Next, we found that the protein A-, but not C- or D-, bands arising from basal Nrf1α in HepG2 cells disappeared to be undetected following *in vitro* deglycosylation reactions with Endo H (Figure S2B, *left two lanes*), implying that protein-A form is a full-length glycoprotein of Nrf1α. This is supported by additional data obtained from LDS-NuPAGE (containing 4-12% polyacrymide,) to separate proteins from total lysates of MG132 (1 μmol/L)-treated cells (Figure S2D). The results revealed that deglycosylation with Endo H enabled for digestion of Nrf1α glycoprotein to disappear, but other three derivative forms appeared to be unaffected. Intriguingly, a parallel experiment using the routine SDS-PAGE (containing 8-12% polyacrymide) showed that deglycosylation reactions with Endo H led to a decrease, but not an abolishment, in the MG132-stimulated abundance of Nrf1α glycoprotein A-form (Figure S2B, *right two lanes*). Albeit such a difference in the glycoprotein migration on between LDS-NuPAGE and SDS-PAGE gels remains to be determined in detail, it suggests other not-yet-identified post-translational modifications leading to distinct folding conformations of Nrf1/TCF11, in particular proximity to its N-glycosylation sites, under MG132-stimulated conditions.

### Dose-dependent effects of proteasomal inhibitors on abundances of Nrf1 isoforms with distinct behaviours

By comparison with distinct profiling of Nrf1α/TCF11 in different cell lines as shown previously [48], basal expression of all the endogenous proteins and derivative isoforms were determined at considerable lower levels to be hardly seen (Figure 1F to H). Surprisingly, two closer protein C/D-forms were obviously increased following treatment of HepG2 cells with lower doses of proteasomal inhibitors BTZ (at 0.01 μmol/L), MG132 (at 1 μmol/L) and expoxomicin (Epox, at 0.01 μmol/L). Amongst these three chemicals, the lowest dose of Epox also caused an increase in the another two longer protein A/B-forms (Figure 1,G & H). Concomitantly with the increasing doses of Epox from 0.01 to 10 μmol/L, abundances of the protein A/B-forms were modestly enhanced, but this was accompanied by gradual decreases in the protein C/D-forms. In the parallel experiments, similar dose-effect relationships were also determined in other two cases of BTZ and MG132 (Figures 1G,H, and S2E). However, it is important to note that an increase in the protein C/D-form accumulation induced by BTZ or Epox at a low dose (at 0.01 μmol/L) was only modestly reduced, but not abolished, by a high dose (at 10 μmol/L) of MG132 (*lanes 11*,*12*). The pattern alternation was not accompanied by a resultant enhancement in abundance of the protein A/B-form, even though the protein C-form seemed to disappear. Overall, our experimental evidence do not support the previous notion reported by the authors referred to [37].

To further clarify dose-dependent effects of proteasomal inhibitors on distinct Nrf1α/TCF11 isoforms, similar cell lysates were subjected to protein separation by 4-12% LDS-NuPAGE and visualized by Western blotting with three different antibodies against distinct regions of Nrf1 (i.e. D5B10 from Cell Signaling Technology, H285 from Santa Cruz Biotechnology, besides one raised in the Zhang’s own laboratory). Together with comparison to ectopically V5-tagged mNrf1 (Figures 1B,C & S2C,D), endogenous Nrf1α glycoprotein was migrated at ^~^120-kDa estimated on LDS-NuPAGE gels (Figure 1H), although it was resolved by routine SDS-PAGE gels to be the protein-A of ^~^140-kDa as estimated above. The glycoprotein should be unstable to be rapidly degraded or processed into other short isoforms, because it was faint to be hardly detected under untreated conditions, but also not substantially increased even upon stimulation by BTZ, Epox and MG132. By contrast, relative accumulation of the ^~^95-kDa full-deglycoprotein (which was resolved by SDS-PAGE gels to be the protein-B/B^−^ form on some occasions, when it was likely mixed with a small fraction of partially-deglycoprotein and/or the N-terminally-truncated glycoprotein between ^~^105-kDa and 110-kDa estimated) was significantly stimulated by proteasomal inhibitors, and the stimulated abundance was gradually increased with the increasing dose of these proteasomal inhibitors (Figure 1, H2 & H3). This was coincidently accompanied by gradual decreases in the protein C/D-forms, that were migrated to between 90-kDa and 85-kDa during NuPAGE, as the doses of proteasomal inhibitors increased. This suggests that both protein C/D-forms should be generated from putative processing of the protein A/B-forms. Moreover, additional processed isoform of ^~^75-kDa (that was only recognized by antibodies against Nrf1β, but not by other two Nrf1 antibodies H285 and D5B10 as described above) was significantly increased with increasing doses of proteasomal inhibitors (Figure 1H1), implying that it should be an N-terminally AD1-truncated isoform. Curiously, the abundance of the ^~^75-kDa isoform stimulated by 0.01 μmol/L of Epox appeared to be partially suppressed by 10 μmol/L of MG132 (Figure 1H1, *cf. lanes 4*,*5 with 11*), as consistent with the previous notion reported by the authors referred to [37]. All together, Nrf1α/TCF11 is allowed for successive glycosylation, ensuing deglycosylation, proteolytic cleavage and degradation to give rise to multiple isoforms, each with distinct behaviours being migrated during LDS-NuPAGE and SDS-PAGE.

To establish a criterion generally acceptable for the entry to work on endogenous Nrf1α/TCF11 and derivatives, similar dose-effect experiments as described above have been extensively carried out in eight other cell lines. These included four human cell lines of HL-7702 (Figure S2F), SH-SY5Y, MHCC97L and HEK293 (Figure S3, A to C), two rat cell lines of RL-34 and Tendon (Figure S4, A & B), and additional two mouse cell lines JB6 and GC-2 (Figure S4, C & D). In all these cases, three to six isoforms (i.e. polypeptides A- to F-bands) were separated by routine 8-12% SDS-PAGE gels running for distinct lengths of time and visualized by immunoblotting with three different antibodies against Nrf1 (Figures S2 to S4). The tricky variations in their abundances could be dependent on different cell types, in which various amounts of Nrf1α and/or TCF11 were differentially expressed and partially processed, leading to distinct profiling of multiple proteaforms, though they had been under similar basal or proteasomal inhibitors-stimulated conditions. Of note, it was found that there exists TCF11 in the normal human and rat, but not mouse, cell lines [51–53, 63], with it being scarcely expressed in cancer cell lines (which was confirmed by sequencing data not shown). In the neuroblastoma SH-SY5Y cells (Figure S3A), as the doses of MG132, Epox and BTZ increased, abundances of protein-A and/or -C forms of Nrf1 were gradually diminished to their disappearances, then they were replaced by gradually increasing protein-B, along with concomitantly decreasing protein-D isoforms. Similar results were obtained from examinations of other cell lines, such as MHCC97L, HEK293, RL-34, Tendon and GC-2 cells (Figures S3 and S4).

Two other polypeptide isoforms between 110-kDa and 100-kDa (recognized by anti-Nrf1 antibodies from Santa Cruz Inc and Cell Signaling Inc, respectively) were also presented in SH-SY5Y cells (Figure S3, A1 & A3), but generation of the ^~^100-kDa isoform was obviously prevented by Epox only (*lanes 31-33 vs 37*). Treatment of GC-2, RL-34 and Tendon cells with either MG132 or BTZ also caused an inducible accumulation of another ^~^110-kDa protein-E isoform (recognized by antibodies against Nrf1β) in a dose-dependent manner (Figure S4, A & B, *lanes 2 to 11*). Moreover, MG132-stimulated abundance of a ^~^90-kDa isoform (which was only recognized by Nrf1 antibodies from Santa Cruz Inc.) was determined in HEK293 cells alone (Figure S3C1, *lanes 3 to 5*). In addition, distinct dose-dependent effects on all four Nrf1α-derived isoforms A to D were also caused by calpain inhibitor-I (CI, also called ALLN, acting as a proteasomal co-inhibitor [46]), but not by CII or calpeptin (CP) (Figure S4E). This indicates that putative proteolytic processing of endogenous Nrf1α/TCF11 is unaffected by calpains.

### Time-dependent effects of proteasomal inhibitors on accumulation of Nrf1 isoforms with distinct stabilities

To elucidate the order of putative post-translational modification events whereby Nrf1α/TCF11 is processed to yield multiple isoforms, here we have carried out time-course analyses of distinct dose-dependent effects of proteasomal inhibitors on the CNC-bZIP protein and derivatives. The results revealed that, with the increasing time from 2- to 24-h of exposure of HepG2 cells to a lower dose of either MG132 (at 1 μmol/L) or BTZ (at 0.01 μmol/L), all four major Nrf1α-derived isoforms A to D were increased before terminating the pulse-chase experiments (Figure 2, A & B, *lanes 1-6*), but the order of such increased isoforms was inverted (from D to A) as graphically shown (*lower left panels*). This convincingly demonstrates that such time-dependent increases in abundances of the major protein-D and minor protein-C isoforms are stimulated by lower concentrations of proteasomal inhibitors, even though with a lag modest enhancement in the other longer A/B-isoform being also induced. By contrast, the higher dose-effects of MG132 (at 10 μmol/L) or BTZ (at 10 μmol/L) were also exerted, as a consequence, leading to a dominantly-stimulated increase in the longer protein-B isoform, which was also accompanied by minor lagged increases in the intact glycoprotein-A form of Nrf1α and processed isoform-D (Figure 2A & B, *lower right graphs*). Similar results were obtained from other parallel pulse-chase experiments (Figure S5, which were validated by Western blotting with another anti-Nrf1 antibody D5B10 from Cell Signaling Technology). In addition, such a higher dose at 10 μmol/L of BTZ also stimulated a time-dependent increase in abundances of both protein-E and -F forms (Figure 2B, *lanes 7 to 11*).

**Figure 2.**
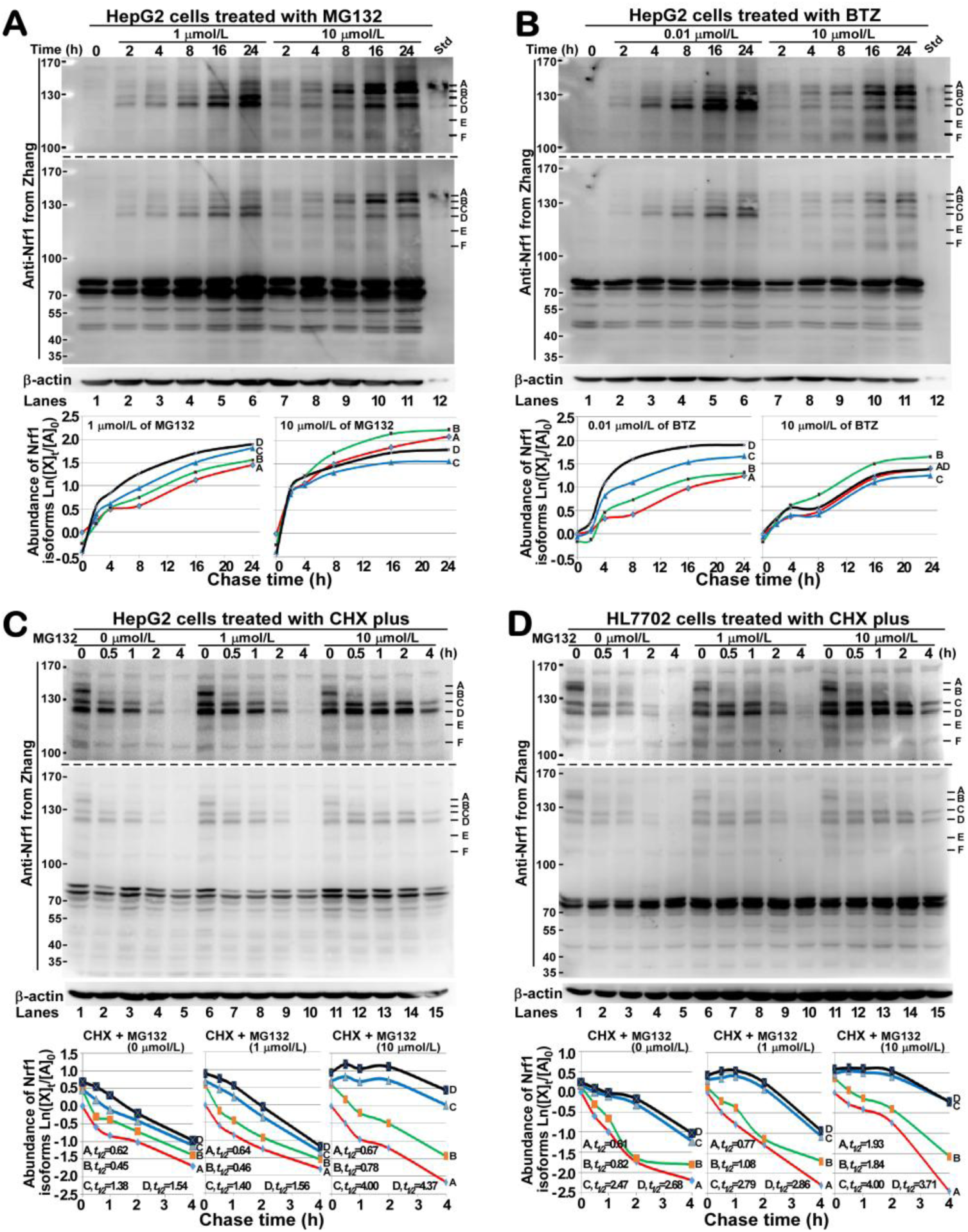
Time-dependent effects of proteasome inhibitors on endogenous Nrf1 isoforms with distinct half-lives. **(A,B)** HepG2 cells were treated with distinct concentrations of MG132 (A) or BTZ (B) for distinct lengths of time (from 0 to 24 h) before being harvested. The total lysates were resolved by SDS-PAGE gels (with upper and lower two-layers containing 8% and 12% polyacrylamide, respectively) and then visualized by immunoblotting with Zhang’s antibody against Nrf1. The relative intensity of four major bands electrophoretically representing distinct Nrf1 isoforms were quantified and shown graphically on the bottoms. **(C, D)** Experimental HepG2 (C) and HL7702 (D) cells were treated with cycloheximide (CHX, 50 μg/ml) alone or plus a lower (1 μmol/L) or higher (10 μmol/L) doses of MG132 for different times before being disrupted. Subsequently, the abundances of distinct Nrf1 isoforms were determined by SDS-PAGE gels with two-layers as described above. The upper images were cropped from the lower pictures, both of which were exposed to the development reagents for different times. The half-lives of four major Nrf1 isoforms were calculated and shown graphically on the bottoms.

Next, to determine the stability of endogenous Nrf1α/TCF11 proteins and derivative isoforms, both HepaG2 and HL-7702 cell lines were treated with 50 μg/ml of cycloheximide (CHX, that inhibits biosynthesis of nascent proteins) alone or together with the proteasomal inhibitor MG132. Western blotting of these lysates showed that abundances of all four Nrf1α/TCF11-derivated protein-A to -D isoforms were significantly decreased following treatment with CHX for 30 min to 2 h, before their disappearance by 4-h treatment (Figure 2C,D, *lanes 1 to 5*). The stoichiometric graphs (Figure 2C, *lower left panels*) showed a tendency to reveal a marked difference in the conversion of between these four Nrf1α-derived A, B, C and D isoforms. Their distinct half-lives were determined to be 0.62 h(=37 min), 0.45 h (=27 min), 1.38 h (=83 min) and 1.54 h (=92 min), respectively, after CHX treatment of HepG2 cells. By comparison with the Nrf1α-expressing HepG2 cells, a nuance in the stability of similar patterned protein-A to -D isoforms arising from co-expression of Nrf1α and TCT11 was found with their distinct half-lives estimated to be 0.61 h (=36.5 min), 0.82 h (=49 min), 2.47 h (=148 min) and 2.68 h (= 160 min), respectively, after treatment of HL-7702 cells with CHX (Figure 2D, *lower left graph*). This indicates that a fraction of each isoforms may be also derived from putative processing of endogenous TCF11 in addition to Nrf1α proteins.

When compared with the above results from treatment with CHX alone, the half-lives of distinct Nrf1α/TCF11-derived isoforms (i.e. protein-A to -D) was almost not or only less prolonged, though their inducible abundances were differentially increased, by co-treatment of HepG2 and especially HL-7702 with a lower dose (at 1 μmol/L) of MG132 for 30 min to 4 h (Figure 2, C & D, *lanes 6 to 10*, and *lower middle graphs*). By contrast, abundances of all four isoforms of Nrf1α/TCF11, particularly its processed proteins C- and D-isoform, were markedly increased until 4 h by co-treatment of experimental cells with MG132 at a higher dose (10 μmol/L) (Figure 2D, *lanes 11 to 15*). Consequently, the stability of such inducible protein isoforms A to D was significantly enhanced, so that their distinct half-lives were extended to 1.93 h (=116 min), 1.84 h (=110 min), 4.00 h (=240 min) and 3.71 h (=222 min), respectively, after co-treatment of HL-7702 cells with a higher dose of MG132 (*lower right graph*). Another similar but different enhancement in stability of Nrf1α-derived proteins A to D (Figure 2C, *lanes 11-15*) occurred following MG132 co-treatment of HepG2 cells, which was determined with their half-lives estimated to be 0.67 h (=40 min), 0.78 h (=47 min), 4.00 h (=240 min) and 4.37 h (=262 min), respectively after CHX treatment (*lower right graph*).

### Knockout of Nrf1α, but not Nrf2, prevents dose-dependent induction of Nrf1-target proteasomal subunit genes by a lower rather than higher concentration of proteasomal inhibitors through the feedback response recruit

To determine which dose-effects of proteasomal inhibitors MG132 and BTZ on endogenous Nrf1α-derived factors are exerted to trigger transcriptional expression of Nrf1-target proteasomal subunit genes, two different concentrations of these two chemicals were subjected to treatment of HepG2 cells that had been transfected with ARE-driven gene reporters (i.e. *GSTA2-6×ARE-Luc* and *3×PSMA4-Luc*, as described elsewhere [11, 35, 46]). Intriguingly, either BTZ or MG132 at the indicated lower and higher doses caused transactivation activity of two reporter genes (Figure 3A), of which incremental induction was notably stimulated by a higher (10 μmol/L), but not lower (i.e. 0.01 or 1 μmol/L), dose of the two inhibitors. Further experiments revealed that the above significant transactivation of reporter genes resulted principally from a remarkable increase in the inducible abundance of endogenous Nrf2 factor (Figure 3B, *left middle panel*). This was concomitantly stimulated by the higher doses of BTZ and MG132, although both chemicals caused an accumulation of ubiquitinated proteins (*right upper panel*). This is supported by the data, as anticipated, revealing that the increased transactivation activity of two ectopic reporters stimulated by higher doses of BTZ and MG132 was substantially repressed by deficiency of Nrf2 to nearly levels obtained from induction by lower doses of the proteasomal inhibitors (Figure 3C). The *Nrf2*^−/−^ cell line, that was established from HepG2 cells with a site-specific loss of the indicated region of the *Nfe2l2* gene manipulated by CRISPR-Cas9-mediated genome editing, was validated by Western blotting analysis (Figure 3D1). Yet, no changes in the abundances of Nrf1α-derived isoforms in between *Nrf2*^−/−^ (Figure 3D3) and its progenitor HepG2 cells (Figure 3B, *left upper panel*) were observed.

**Figure 3.**
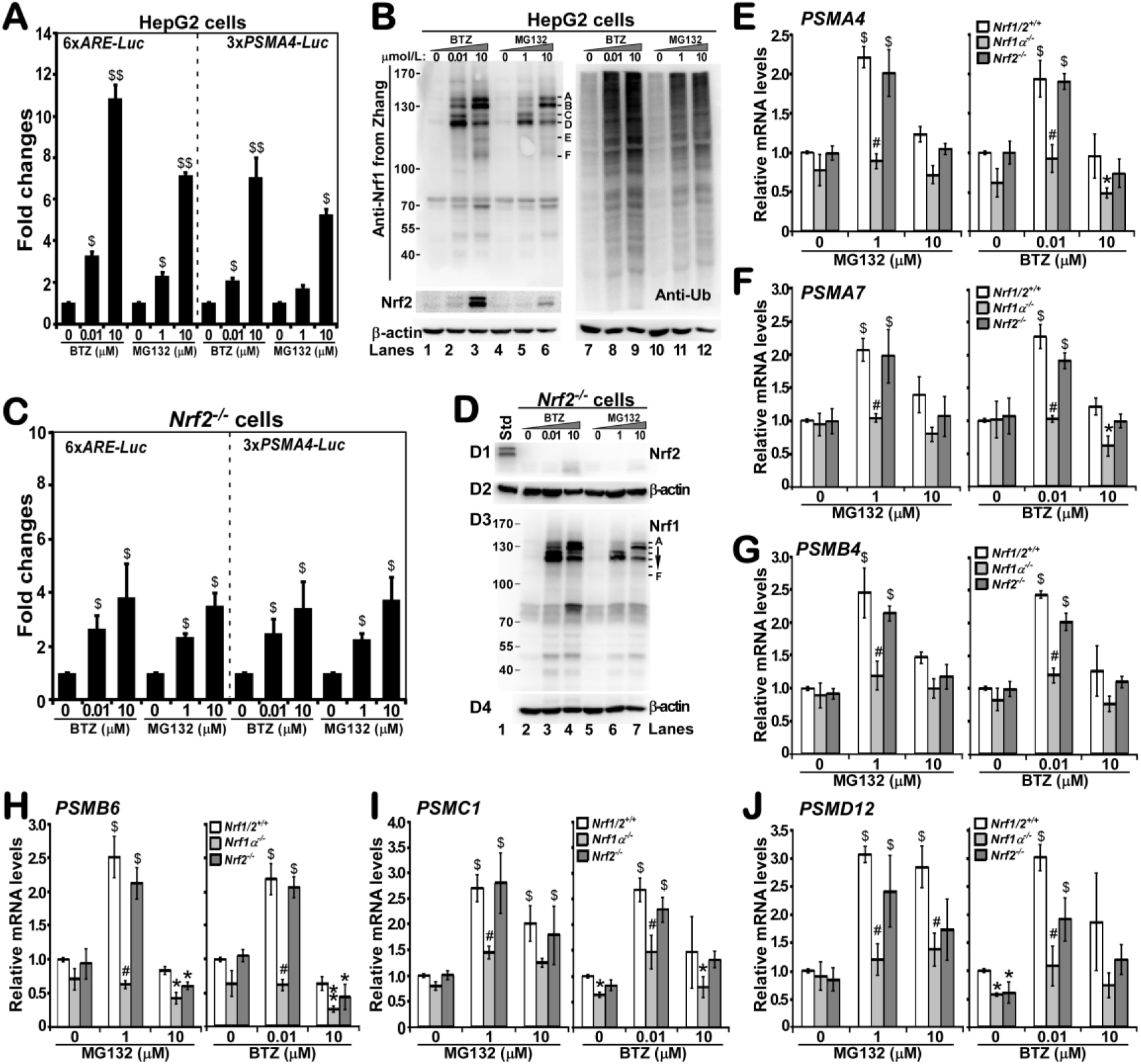
Diverse dose-dependent effects of proteasomal inhibitors on Nrf1/2-mediated gene expression. **(A,B)** HepG2 cells were transfected for 7 h with either *P_SV40_GSTA2-6×ARE-Luc* or *3×PSMA4-ARE-Luc* reporter (120 ng of DNA), along with another *Renilla* reporter (60 ng) as an internal control for transfection efficiency. The cells were allowed for a 16-h recovery from transfection and then treated with distinct doses of proteasomal inhibitors MG132 and BTZ for additional 16 h before the ARE-driven luciferase reporter activity was measured (A). The experimental data were calculated as a fold change (mean ± S.D) relative to the basal activity of untreated group (at the value of 1 given) with significant increases ($, p < 0.05; $$, p < 0.01). Similarly, the proteasomal inhibitors-treated cell lysates were also separated by SDS-PAGE with two layers that consist of 8% and 12% polyacrylamide respectively, and then visualized by immunoblotting with antibodies against Nrf1, Nrf2 or Ub. **(C,D)** *Nrf2*^−/−^-specific knockout (established by CRISP-Cas9-mediated editing of the genome in HepG2) cells were subjected to similar chemical treatments, followed by measurement of ARE-driven reporter activity (C) and visualization by Western blotting (D), as described above. **(E to J)** Relative basal and stimulated expression levels of distinct proteasomal subunit genes (including *PSMA4*, *PSMA7*, *PSMB4*, *PSMB6*, *PSMC1* and *PSMD12*) at their mRNA levels were determined by real-time qPCR analyses of different three cell lines that had been treated with different doses of proteasome inhibitors for 16 h. The resulting data presented each represent at least three independent experiments undertaken on separate occasions, and significant increases was also calculated ($, p < 0.05; $$, p < 0.01).

Next, distinctive dose-dependent effects of proteasomal inhibitors on the endogenous expression of Nrf1-target genes, such as those encoding all the 26S proteasomal subunits [11, 35, 37], were further determined by quantitative real-time PCR analysis of both the wild-type HepG2 (*Nrf1/2^+/+^*) and derived mutant cell lines (*Nrf1α*^−/−^ and *Nrf2*^−/−^). The results (Figure 3,E to J) showed that significant increases in transcriptional expression of all six genes *PSMA4*, *PSMA7*, *PSMB4*, *PSMB6*, *PSMC1* and *PSMD12* were examined following treatment of *Nrf1/2^+/+^* cells with a lower dose (at 0.01 or 1 μmol/L) of BTZ and MG132. Such BTZ/MG132-inducible gene expression was completely abolished by knockout of *Nrf1α*^−/−^ but not of *Nrf2*^−/−^. However, none of all four core genes *PSMA4*, *PSMA7*, *PSMB4 and PSMB6* (which are representatives of the 20S core particle of proteasome) were stimulated by a higher dose (at 10 μmol/L) of BTZ or MG132 (Figure 3, E to H). By contrast, other two regulatory genes *PSMC1* and *PSMD12* (as representatives of the 19S regulatory particle of proteasome) were modestly induced by 10 μmol/L of MG132 rather than BTZ (Figure 3I, J), but their modest induction was also blocked by knockout of *Nrf1α*^−/−^ . Collectively, these demonstrate that Nrf1α-derived isoforms, but not Nrf2, are required especially for mediating transcriptional expression of all examined proteasomal subunit genes, that are induced in the feedback response to stimulation by a lower dose of proteasomal inhibitors.

### Both p97 knockdown and its inhibitor NMS-873 exert similar negative effects on the Nrf1α-related processing

The versatile AAA-ATPase p97 was shown to drive dynamic retrotranslocation of ectopically-expressed Nrf1 from the ER luminal side of membranes into the cyto/nucleoplasmic side, whereupon the CNC-bZIP protein could be allowed for deglycosylation digestion and proteolytic processing to give rise to multiple isoforms [46, 57]. However, none of available evidence has been provided insofar as to demonstrate convincingly that p97-driven retrotranslocation of endogenous Nrf1α/TCF11 enables its intact proteins to be extracted from the ER and then processed on the extra-ER luminal side of membranes [37, 55, 56]. To address this issue, HepG2 cells were allowed for significant knockdown of p97 to considerable lower levels (Figure 4A, *upper panel*) by transfecting three different oligonucleotide sequences of si-p97, i.e. small-interfering RNAs (siRNAs) targeting to distinct coding regions of p97 transcript (numbered as 1, 2 and 3). The results further revealed that knockdown of p97 caused an obvious accumulation of endogenous Nrf1α -derived protein-A to -D isoforms (*middle panel*), when compared with that obtained from a negative (-Ve) control of transfection with a scrambled siRNA.

**Figure 4.**
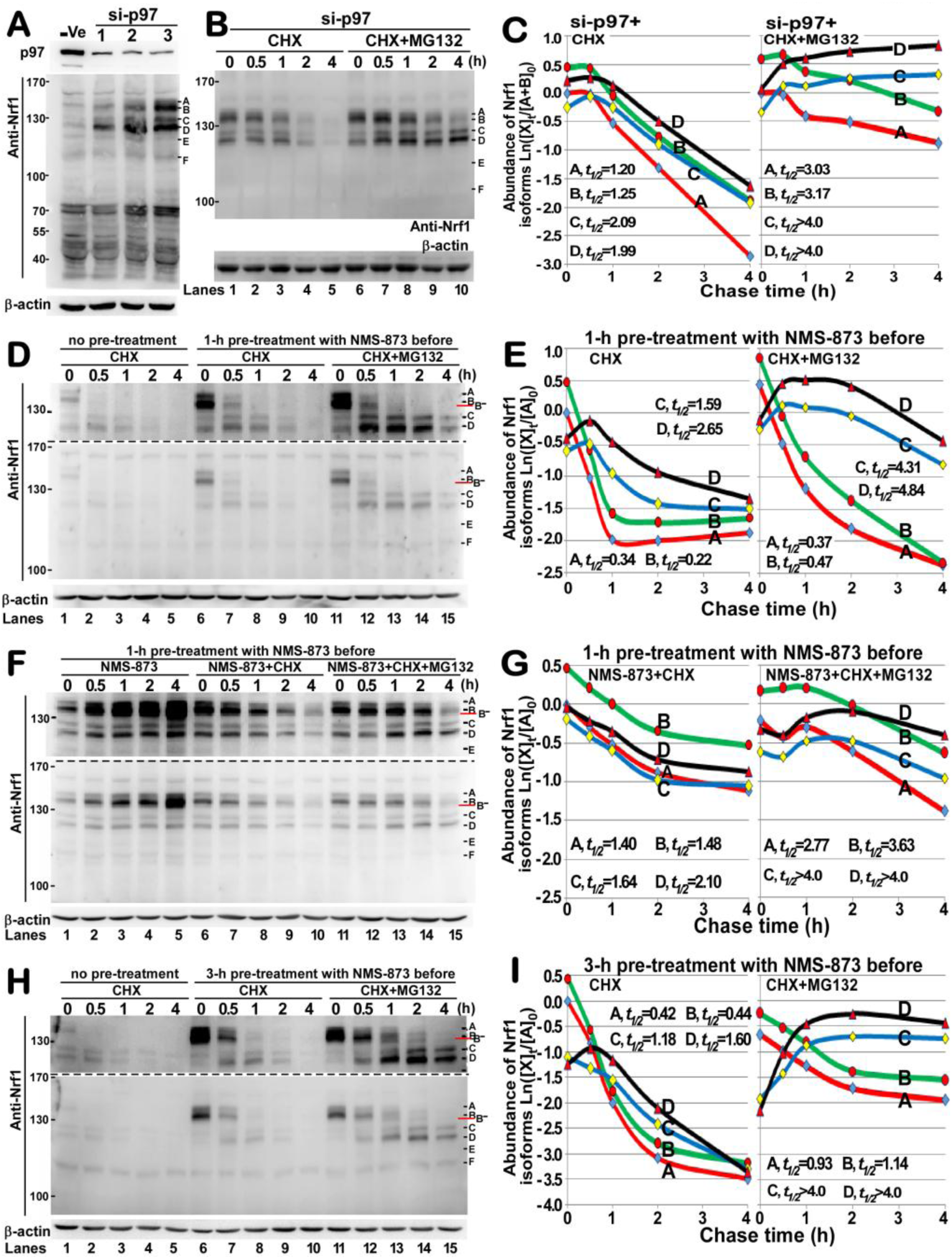
Effects of the p97 AAA-ATPase on the processing of endogenous Nrf1 proteins. **(A)** Total lysates of HepG2 cells that had been transfected with each (50 nmol/L) of three different siRNA sequences targeting against p97 (i.e. si-p97), along with a scrambled siRNA as an internal negative (-Ve) control, were resolved by SDS-PAGE containing 10% polyacrylamide, and then analyzed by Western blotting with antibodies against p97 or Nrf1. **(B,C)** HepG2 cells that had been transfected with the third si-p97 sequence were allowed for a 16-h recovery culture, and then treated with CHX (50 μg/ml) alone or in combination with MG132 (10 μmol/L) for distinct indicated times before being disrupted. Relative abundances of four major Nrf1 isoforms were determined by immunoblotting (B), with their distinct half-lives being calculated as shown graphically (C). **(D to G)** HepG2 cells were or were not pretreated with a p97-specific inhibitor NMS-873 (10 μmol/L) for 1 h, before being changed in another fresh culture medium to which CHX (50 μg/ml) alone or plus MG132 (10 μmol/L) were added, and being harvested at different time points. The total lysates were resolved by SDS-PAGE (with two-layers made of 8% and 12% polyacrylamide, respectively) and then examined by immunoblotting with Zhang’s Nrf1 antibody (D). Subsequently, the intensity of four major Nrf1 protein bands was quantified and shown graphically (E). Similarly, other experiments of HepG2 cells that had been allowed for another 3-h pretreatment with NMS-873 (F,G) were done as described above. **(H,I)** HepG2 cells were pretreated with NMS-873 (10 μmol/L) for 1 h, and then allowed for transfer another fresh culture medium containing DMSO, CHX (50 μg/ml) alone or plus MG132 (10 μmol/L) before being disrupted. Relative expression of major Nrf1 isoforms in the total lysates was determined by Western blotting (H) and also shown graphically (I).

Next, the pulse-chase experiments of HepG2 cells that had been transfected with si-p97 were employed, aiming to determine the order of conversion and/or turnover of distinct Nrf1α-derived isoforms, with different topovectorial behaviours, that were altered following inhibition of putative p97-driven retrotranslotion. As expected, time-course analysis of the No.3 si-p97-transfected HepG2 cells, which was treated with 50 μg/ml of CHX alone for 30 min to 4 h (Figure 4B, *lanes 1-5*), revealed that the stability of endogenous Nrf1α-related protein-A to -D forms was strikingly enhanced (Figure 4C, *left panel*). The resultant half-lives were markedly extended to 1.20 h (=72 min), 1.25 h (=75 min), 2.09 (=125 min) and 1.99 h (=119 min) respectively, after CHX treatment, when compared with those measured from the untransfected cells (i.e. 0.62, 0.45, 1.38 and 1.54 h, Figure 2C, *lower left panel*). By contrast, co-treatment of si-p97-transfected cells with CHX and MG132 (at 10 μmol/L) further caused a profound increase in the stability of distinct Nrf1α-derived protein-A to D isoforms (Figure 4B, *lanes 6-10*), such that their turnover was predominantly prolonged as shown graphically (Figure 4C, *right panel*). As a matter of fact, the result led to significant prolongation of distinct half-lives of these Nrf1α-derived isoforms; i.e. protein-A and -B were estimated to extend to 3.03 h (=182 min) and 3.17 h (=190 min), whilst both protein-C and D were defined to be over 4 h (=360 min), respectively.

Intriguingly, similar but different results were obtained from additional pulse-chase experiments of HepG2 cells that had been treated with the p97-specific inhibitor NMS-873 (at a dose of 10 μmol/L). As shown in Figure 4D, dominant increases in abundances of endogenous Nrf1α-derived protein-A and -B, as accompanied by only minor increases in both protein-C and -D isoforms, were observed in the cells that had been pre-treated with NMS-873 for 1 h before being treated with CHX for indicated times (*lanes 6-10*), when compared with the data obtained from the cells that had been not pre-treated with NMS-873 before addition of CHX (*lanes 1-5*). Curiously, such inhibition of p97-dirven retrotranslocation by NMS-873 and subsequent release from the inhibitor were allowed for time-course analysis of the existing Nrf1 protein-A to -D conversion in order. The resultant alteration in these protein stability was determined with distinct half-lives estimated to be 0.34 h (=20 min), 0.22 h (=13 min), 1.59 h (=95 min) and 2.65 h (=159 min) after CHX treatment, respectively (Figure 4E, *left panel*), implying a time-dependent order of conversion of Nrf1α-derived protein-A to -D isoforms. However, co-treatment of NMS-873-pretreated cells with CHX and MG132 did not cause a further increase in abundances of Nrf1 protein-A and -B isoforms (Figure 4D, *lanes 11-15*). Also, no remarkable changes in their half-lives to disappear were observed, that were estimated to be 0.37 h (=22 min) and 0.47 h (=28 min) respectively (Figure 4E, *right panel*). By contrast, the major protein-D, alongside with the minor protein-C, isoforms (arising from the existing Nrf1 protein-A and -B isoforms) were accumulated from 30 min to 2 h following co-treatment of NMS-873-pretreated cells with CHX and MG132 (Figure 4D, *lanes 11-15*). Thus, turnover of protein-C and -D was significantly postponed by the proteasomal inhibitor, that was determined with their half-lives prolonged to 4.31 h (=258 min) and 4.84 h (=290 min), respectively (Figure 4E, *right panel*).

While the time of treatment of the 1-h NMS-873-pretreated cells with the same chemical was increasing from 0.5 to 4 h, the resulting extension of p97 inhibition led to a gradual incremental accumulation of Nrf1α-related minor protein-A and major protein-B isoforms (Figure 4F, *lanes 1-5*). Additional 3-h NMS-873 pretreatment of HepG2 cells also caused a substantial increase in abundances of Nrf1α-related protein-A and -B, when compared to both levels measured from no pretreatment of cells with NMS-873 (Figure 4H, *lanes 6 vs 1*). Further pulse-chase experiments with addition of CHX revealed that the disappearance of the existing Nrf1α-related protein-A and -B isoforms was delayed by 3-h pretreatment with NMS-873 (*lanes 6-10*), with their half-lives extended to be 0.42 h (=25 min) and 0.44 (=26 min) (Figure 4I, *left panel*), relative to equivalents obtained following 1-h pretreatment with this inhibitor (i.e. 0.34 and 0.22 h, Figure 4E, *left panel*). By contrast, the stability of minor protein-C and major protein-D isoforms, with respective half-lives of 1.18 h (=71 min) and 1.60 h (=96 min) (Figures 4I, *left panels*), was obviously reduced by 3-h pretreatment with NMS-873, relatively to equivalents determined following 1-h pretreatment with the inhibitor (i.e. 1.59 and 2.65 h, Figure 4E, *left panel*). These demonstrate that the extent of p97-arrested retrotranslocation of intact Nrf1 from the luminal side of membranes into extra-ER compartments monitors the ordered processing of the CNC-bZIP protein to yield multiple isoforms. This is also supported by further time-course analysis of co-treatment of HepG2 cells with CHX and MG132 following 3-h pre-treatment of NMS-873. The results indicated that the stability of all these Nrf1α-derived protein-A to -D isoforms was significantly enhanced by the proteasomal inhibitor (Figure 4H, *lanes 11-15*), with their prolonged half-lives as shown graphically in (Figure 4I, *right panel*).

Further pulse-chase experiments unraveled that distinctive time-dependent effects of NMS-873 on Nrf1α-derived isoforms A to D (Figure 4F, *lanes 6-10*) were elicited to postpone conversion of these existing CNC-bZIP proteins, in particular longer isoform-A and B. This was determined with half-lives of protein-A and -B prolonged to 1.40 h (=84 min) and 1.48 h (=89 min), as accompanied by no obvious changes in half-lives of protein-C and -D estimated to be 1.64 h (=98 min) and 2.10 (=126 min) respectively, following co-treatment with CHX (Figure 4G, *left panel*). Moreover, the stability of these proteoforms was modestly promoted by addition of MG132 into the cultured medium of cells that had been pre-treated for 1 h with NMS-873 and then continued to co-treat with CHX for indicated times (Figure 4F, *lanes 11-15*). The co-treatment enabled the half-lives of two longer protein-A and -B to be profoundly prolonged to 2.77 h (=166 min) and 3.63 h (=218 min), respectively (Figure 4G, *right panel*); this was also accompanied by an extension of half-lives of protein-C and -D to over 4 h. All together, dynamic retrotranslocation of Nrf1α, driven by p97-dependent machineries, dictates topovectorial repositioning of the CNC-bZIP protein from the ER luminal side of membranes into the extra-ER cyto/nucleoplasmic side, in which it may be degraded by proteasomes and/or cleaved by other proteases in order to yield multiple isoforms.

### Knockdown of Hrd1 causes an accumulation of the longer isoforms of Nrf1α with the unaltered turnover

To the best of our current knowledge [11, 36, 64–67], it is inferable that the membrane-bound Nrf1α is ubiquitinated by Hrd1, an ER membrane-anchored ubiquitin ligase, that may trigger retrotranslocation of a large luminiar-resident portion of the CNC-bZIP protein from the luminal side across membrane lipid bilayers into the cyto/nucleoplasmic side. This topovectorial process is driven by a direct Hrd1-interactor p97, that enables its client protein Nrf1α to be unleashed from the membrane confinements so as to dislocate into extra-ER subcellular compartments, prior to its mature processing into multiple isoforsms. Conversely, knockdown of Hrd1 by si-Hrd1 (i.e. a small-interfering RNA targeting to Hrd1, Figure 5A) led to an accumulation of endogenous Nrf1, particularly longer protein-A and -B forms in HepG2 cells (Figure 5B, *lanes 2 vs 1*). However, no obvious changes in the conversion of Nrf1α-derived protein-A to -D resulted from si-Hrd1 knockdown (Figure 5B, *lanes 2-6*). The stability of protein-A and -B was unaltered by si-Hrd1, together with almost no changes in both half-lives estimated to be 0.50 h (=30 min) and 0.49 h (=29 min). As such, the turnover of protein-C and -D was modestly accelerated with their half-lives shortened to 1.00 h (=60 min) and 1.16 h (=70 min) respectively, after CHX treatment (Figure 5C, *left panel*), when compared with equivalents measured from the cells without si-Hrd1 knockdown (i.e. 0.62, 0.45, 1.38 and 1.54 h, Figure 2C).

**Figure 5.**
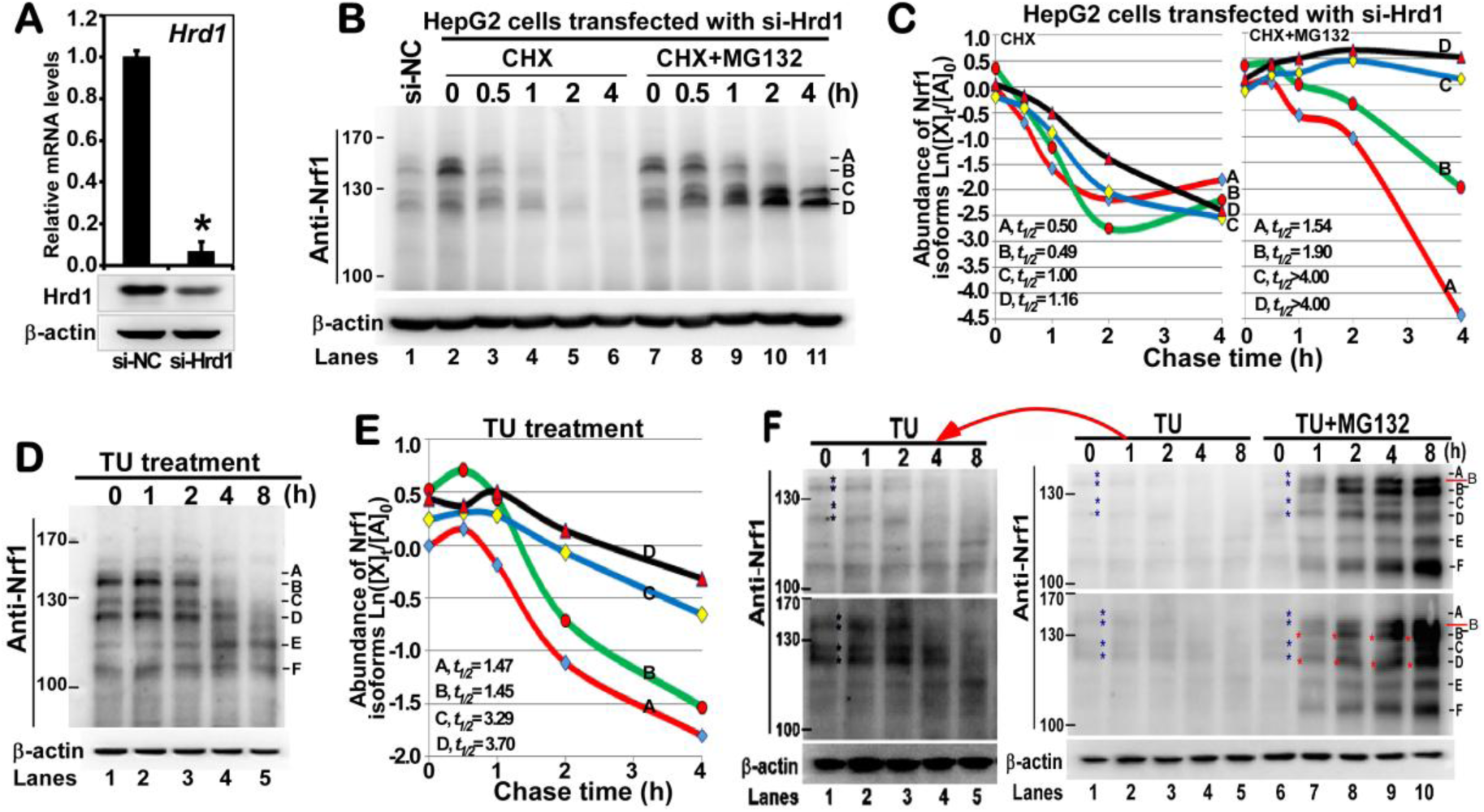
The processing of endogenous Nrf1 monitored by its glycosylation status and Hrd1-mediated events. **(A)** HepG2 cells that had been transfected for 7 h by the lipofectamine 3000 reagent mixed with si-Hrd1, were then allowed for a 24-h recovery culture, and analyzed by real-time qPCR and Western blotting as described for legends of Figure 4. The data obtained from qPCR were calculated as a fold change (mean ± S.D), with a significant decrease (^∗^P < 0.05, n = 3) resulted from si-Hrd1 knockdown, relative to the basal expression of Hrd1 (at the given value of 1, that was measured from siNC-transfected cells). **(B,C)** Similarly transfected HepG2 cells were treated with CHX (50 μg/ml) alone or plus MG132 (10 μmol/L) for indicated times before being lysised in denature buffer. The total lysates were determined by immunoblotting with Zhang’s Nrf1 antibody (B). Relative expression of major Nrf1 protein bands was quantified and calculated as an Ln(X) function value on their fold changes (C). **(D to F)** HepG2 cells were treated with TU (2 μg/ml) alone or in combination with MG132 (10 μmol/L) for indicated times before being disrupted, followed by examination by Western blotting. The results obtained from two independent experiments were presented herein (D,F). Relative abundances of major Nrf1 isoforms were calculated and presented by the Ln(X) function curves (E).

By sharp contrast, further pulse-chase experiments revealed that the stability of Nrf1α-derived protein-A to -D was significantly enhanced upon the proteasomal inhibition by MG132 (at 10 μmol/L) (Figure 5B, *lanes 7-11*), so that their half-lives were substantially prolonged to 1.54 h (=92 min), 1.90 h (=114 min), over 4 h (=240 min) and more than 4 h (=240 min), respectively, after co-treatment of si-Hrd1 cells with CHX (Figure 5C, *right panel*). Together with the previous reports of Hrd1 acting as a retrotranslocon candidate involved in ERAD [64–67], these results indicate that dynamic repositioning of Nrf1α may be partially retarded by knockdown of Hrd1, so that its longer protein-A/B isoforms are modestly accumulated. However, it cannot be ruled out that a small fraction of Nrf1 are permitted by a hitherto unidentified mechanism to be presented for the proteolytic processing of the CNC-bZIP protein by MG132-sensitive proteases, besides proteasomes.

### Post-synthetic modifications of endogenous Nrf1α by glycosylation, deglycosylation and putative cleavage

In fact, it is notable that a few of previous studies by us and other groups [11, 38, 42, 57] had identified that ectopic Nrf1α is N-glycosylated to form an inactive glycoprotein in the lumen of ER. If required, it is then retrotranslocated into the cyto/nucleoplasmic side of membranes, in which the glycoprotein is allowed for deglycosylation digestion to become another active isoform. Both the full-length Nrf1α glycoproteins and deglycoproteins are subject to further proteolytic processing to yield multiple proteoforms. However, no convincing evidence has been presented to date, showing whether endogenous Nrf1α/TCF11 is glycosylated and which isoforms are represented by its glycoprotein, deglycoprotein and truncated proteins, respectively. As such being the case, it is of importance to determine which isoforms of Nrf1α will be firstly influenced during pulse-chase experiments with tunicamycin (TU, a specific inhibitor of oligosaccharyltransferases, and also a classic ER stressor) to block N-linked glycosylation of nascent proteins. For this end, thus we carried out a thorough time-course analysis of TU-treated cells, in order to examine effects of this inhibitor on Nrf1α-derived protein-A to -D isoforms. As anticipated, the results revealed that most of Nrf1 protein-A and -B disappeared after 4 h of treatment of HepG2 cells with TU (2 μg/ml), whereas another lag disappearance of protein-C and -D also occurred by 8 h of this chemical treatment (Figure 5D). The stability of these Nrf1α-derived protein-A to -D isoforms was determined by their different half-lives estimated to be 1.47 h (=88 min), 1.45 h (=87 min), 3.29 h (=198 min) and 3.70 (=222 min), respectively, following TU treatment (Figure 5E).

Upon the proteasomal inhibition by MG132, abundances of all other isoforms of Nrf1 protein-B to -F, except protein-A, were significantly enhanced in a time-dependent fashion (Figure 5F, *lanes 6-10*). However, careful insights into the electrophoretic mobility of Nrf1 isoforms revealed that inhibition of N-glycosylation by TU caused a large majority of its intact protein-A to disappear, but be replaced by gradually enhanced protein-B and/or -B^−^ isoforms (of between ^~^130-kDa and^~^135-kDa) after 2-h to 8-h co-treatment of cells with MG132 (Figure 5F). Interestingly, the disappearance of protein-A was also coincidently accompanied by slightly faster migration of protein-B^−^, -D and -F isoforms, each with a time-dependent increment in the mobility. Collectively, these data indicate co-treatment of MG132 and TU results in a time-dependent accumulation of de-glycosylated (partially and fully), un-glycosylated, and progressively truncated isoforms of Nrf1. Thus, it is postulated that the N-terminal and/or its C-terminal fragments of Nrf1 are much likely to be successively proteolytically removed from this CNC-bZIP protein by other proteasome-independent pathways (e.g. DDI-1/2), whereas the remaining portion of Nrf1 is also unstable to be rapidly degraded by proteasomes. This assumption is further supported by the parallel experimental data (Figure 5F, *lanes 1-5*).

### Post-translational processing of tetracycline-inducible Nrf1α and its derived isoforms by glycosylation, deglycosylation and putative protealysis

To further determine distinct effects of TU alone or plus MG132 on precise Nrf1α-derived isoforms, we established a stable tetracycline-inducible Nrf1α-expressing cell line (i.e. HEK293C^Nrf1α^, which was created on the base of HEK293). Firstly, HEK293C^Nrf1α^ cells were cultured for 12 h in a higher glucose DMEM containing 1 μg/ml of tetracycline to induce stable expression of Nrf1α and its derivatives, before being treated with TU alone or in combination with proteasomal inhibitors MG132 or BTZ. Upon blockage of N-glycosylatio, almost all of major protein-A and minor protein-B of Nrf1 (both were tagged C-terminally by V5 ectope) were completely abolished to disappear within 30 min following treatment with TU (at 2 μg/ml) (Figure 6A, *lanes 1-5*). The disappearance of protein-A/B was, instead, accompanied by a marginal enhancement in other two close protein-C/D isoforms between 30 min and 1 h after TU treatment. This finding was further validated by additional experimental data as illustrated in (Figure 6B, *lanes 6-10*). The stability of major protein-A and -D of Nrf1 was defined by both distinct half-lives estimated to be 0.65 h (=39 min) and 2.91 h (=175 min), respectively, after TU treatment (Figure 6C, *right graph*). By comparison with protein-A and -D, the minor protein-B and -C isoforms were very fainter to be hardly distinguished following over 2-h electrophoresis (Figure 6B). These data indicate a possibility that Nrf1α-derived protein-B and -C may be two transient forms so as to be rapidly converted into other processed protein-D (and -E). In addition, the abundance of Nrf1 processed protein-E was obviously enhanced by treatment with TU, but not TG (thapsigargin) (Figure 6B).

**Figure 6.**
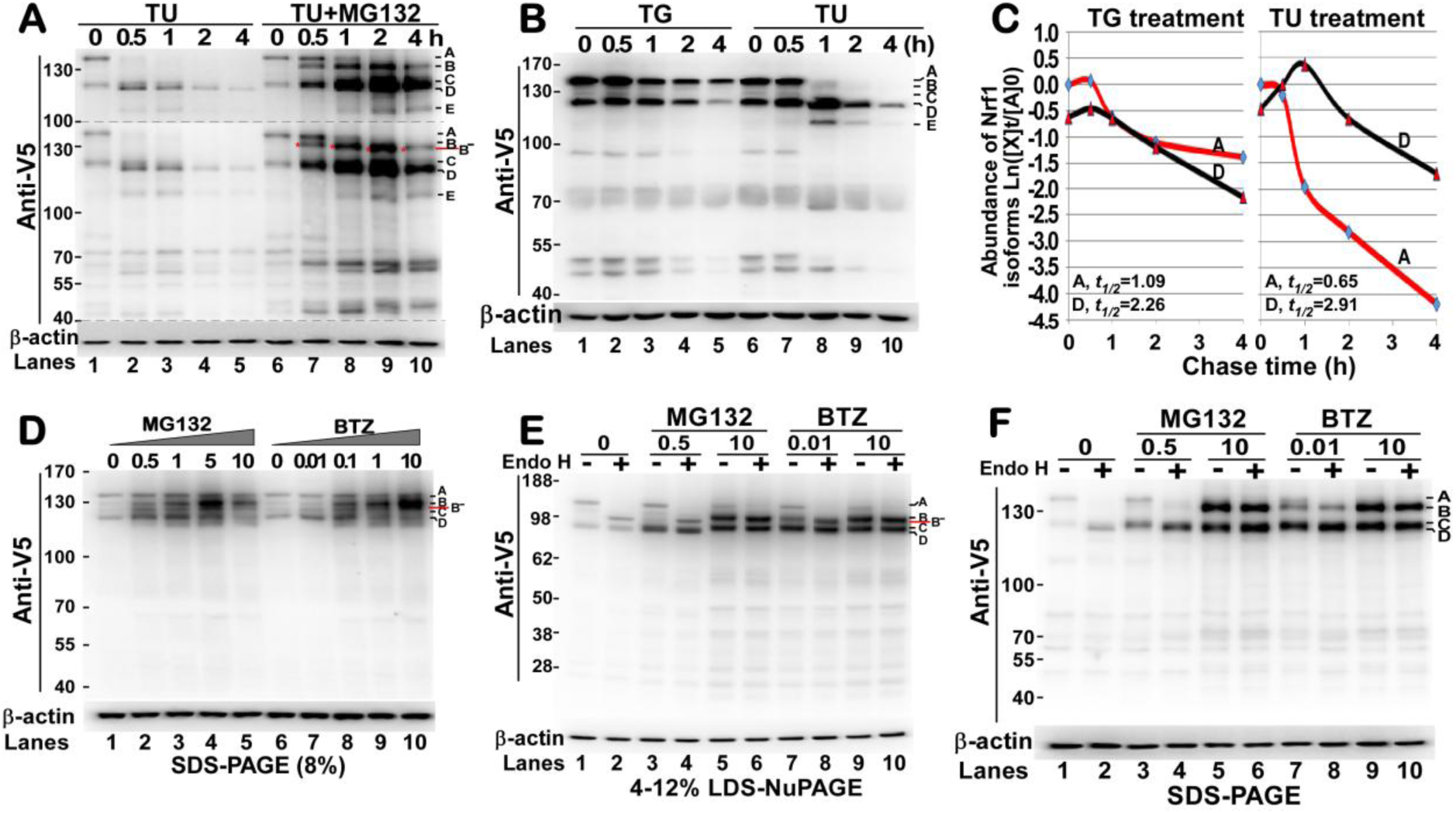
The processing of tetracycline-inducible Nrf1 monitored by its glycosylation status and proteasome-mediated events. **(G to I)** Further pulse-chase experiments were performed in stably tetracycline-induced hNrf1α-expressing HEK293C^Nrf1α^ cells, that had been cultured in the DMEM high glucose medium containing tetracycline (1 μg/ml) for 12 h, before addition of TU (2 μg/ml), TG (1 μmol/L) alone or plus MG132 (10 μmol/L). **(J to L)** HEK293C^Nrf1α^ cells were pretreated with tetracycline (1 μg/ml) for 12 h and then changed for additional 4-h culture in the fresh medium to which different doses of MG132 or BTZ were added. Subsequent lysates were examined by Western blotting with Zhang’s Nrf1 antibodies, and the intensity of its major protein blots, as well as the latter ratios were shown on the bottom (J). Similar lysates were also subjected to *in vitro* deglycosylation reactions with Endo H, followed by protein separation by SDS-PAGE gels with tow-layers consisting of 8% and 12% polyacrylamide (K) or gradient NuPAGE gels containing 4-12% polyacrylamide (L), and visualization by immunoblotting with antibodies against the V5 ectope or β-actin.

By contrast with TU treatment, proteasomal inhibition by MG132 caused a marked increment in the abundances of all Nrf1α-derived protein-A to -E isoforms (Figure 6A, *lanes 6-10*). Amongst these isoforms of Nrf1, most of the protein-A disappeared within 1 h of TU treatment and then seemed to be replaced by incremented protein-B and/or -B^−^ isoforms (Figure 6A, *upper two panels*). The disappearance of protein-A appeared to be accompanied by gradual increases in two close faster-migrated protein-C/D, as well as processed protein-E, within 2 h of co-treatment of cells with TU and MG132. However, all these incremented protein-B to -E isoforms were coincidently allowed to fall into a decline after 4 h of co-treatment with TU and MG132. Similar remarked suppression or abolishment of Nrf1-derived protein-D to -E isoforms also occurred after 2-h treatment of TU alone (Figure 6B, *lanes 8-10*).

By comparison with TU, another classic ER stressor TG exerted time-dependent effects on Nrf1α-derived isoforms (Figure 6B, *lanes 1-5*), which were significantly distinctive from those obtained from TU-treated cells (*lanes 6-10*). The results showed that almost no obvious increases in abundances of Nrf1-derived protein-A to -D were detected within 1 h of TG (1 μmol/L) treatment of HEK293C^Nrf1α^ and thereafter, these isoforms were gradually decreased to lower levels by 4 h of the chemical treatment (Figure 6B). The stability of both major protein-A and -D isoforms was defined by both half-lives estimated to be 1.09 h (=65 min) and 2.26 h (=136 min) after TG treatment (Figure 6C, *left panel*).

These data, together with the previous notion that TG, as a classic ER stressor, can induce the unfolded protein response (UPR) [43, 68], indicate that the intact ER-localized Nrf1 and its derivate isoforms are also influenced by TG-induced ER stress responses, albeit the detailed mechanism remains to be explored.

### Distinct dose- and time-dependent effects of proteasomal inhibitors on the proteolytic processing of tetracycline-inducible Nrf1α into multiple isoforms

To clarify whether the above faster-migrated isoforms of Nrf1 are *de facto* altered by proteasomal inhibitors (or only by it in a combination with TU), here we examined both dose- and time-dependent effects of MG132 or BTZ on Nrf1-derived isoforms in HEK293C^Nrf1α^ cells (Figures 6D and S6). Of note, only two major Nrf1 protein-A and -D were expressed exclusively in the MG132/BTZ-untreated HEK293C^Nrf1α^ cells (Figure 6D, *lanes 1 and 6*). However, abundance of Nrf1 protein-A was only marginally increased by lower doses of MG132 (at 0.5-1.0 μmol/L) or BTZ (at 0.01-1.0 μmol/L, but the marginal effect was not further altered or even repressed by higher doses of MG132 (at 5-10 μmol/L) or BTZ (at 1-10 μmol/L). By contrast, Nrf1 protein-D abundance was markedly increased by lower doses of MG132 (*lanes 2*,*3*) or BTZ (*lanes 7*,*8*), but then decreased or abolished by higher doses of MG132 (*lanes 5*,*6*) or BTZ (*lanes 7*,*8*). Intriguingly, the emergence of Nrf1 protein-C isoform followed stimulation of HEK293C^Nrf1α^ by lower doses of MG132, but it also disappeared due to being prevented by higher doses of this chemical (Figure 6D, *lanes 2-5 vs 1*). Importantly, stimulation of cells by MG132 (*lanes 2-4*) or BTZ (*lanes 8-10*) triggered the emergence of Nrf1 protein-B and resulted in a dose-dependent increase in its abundance. Collectively, these comparisons revealed that there exists an intrinsic significant correlation amongst Nrf1α-derived isoforms and their ordered conversion of protein-A, -B, -C to -D is affected by distinct doses of proteasomal inhibitors.

The above notion is further supported by pulse chase experiments of HEK293C^Nrf1α^ cells with a lower (1 μmol/L) or higher (10 μmol/L) concentration of MG132 (Figure S6). When compared with untreated cells expressing a major protein-A and another minor protein-D (Figure S6A, *left 7 lanes*), the protein-D abundance was significantly enhanced after 1-h treatment with, which was accompanied by stimulated accumulation of protein-B, with a peak of increases at 4 h following treatment and subsequent decreases in their abundances (Figure S6, A to F). Of note, the intact protein-A became gradually fainter to disappear after 4-h treatment with MG132, and thereafter was replaced by enhanced abundance of protein-B, particularly during treatment with 10 μmol/L of MG132. In addition, it should be noted that some experimental differences between HepG2 and HEK293C^Nrf1α^ may also be resulted from removal of tetracycline, that induces transcription of Nrf1, during treatment of HEK293C^Nrf1α^ cells.

The *in vitro* deglycosylation reaction with Endo H revealed that Nrf1 protein-A isoform (i.e. ^~^120-kDa resolved by LDS-NuPAGE) was completely digested by this glycosidase (Figure 6E). Following disappearance of protein-A, it was replaced by the faster-migrated protein-B^−^ (i.e. ^~^95-kDa, *lanes 1-4*), and the abundance of another slower-migrated protein-B (i.e. ^~^105-kDa) was also enhanced with no changes in its mobility (*lanes 5-10*). By contrast, the protein-D (i.e. ^~^85-kDa) was ma accumulated by 0.5 μmol/L MG132 (*lanes 3*,*4 vs 1*,*2*), but no changes in its migration were observed following deglycosylation digestion. Intriguingly, a little shift of protein-B^−^ and -D to the slower locations of protein-B and -C (i.e. ^~^90-kDa) respectively, occurred following treatment with 10 μmol/L MG132 or 0.01-10μmol/L BTZ (Figure 6E, *lanes 5-10*). Further examinations of similar lysates by routine SDS-PAGE showed distinctive patterns of Nrf1α-derived protein-A to -D varying with different conditions (Figure 6F). The protein-A was completely digested by Endo H to disappear, but both protein-B and -D were unaffected by the glycosidase. Collectively, these results indicate that, besides glycoprotein-A (120-kDa), deglycoprotein-B^−^ (95-kDa) and its N-terminally-truncated protein-D (85-kDa), both of protein-B (105-kDa) and protein-C (90-kDa) should be two transient unstable isoforms to be rapidly processed into deglycoprotein-B^−^ and cleaved protein-D, respectively. It is also inferable that the protein-B isoform might be a partial-deglycoprotein or truncated glycoprotein arising from glycoprotein-A, whereas the protein-C may be an insufficiently-truncated form. In addition, a little slower migration of both the 95-kDa and 85-kDa proteins was, though unaffected by degycosylation digestion, also observed following electrophoresis of cell lysates that had been treated with MG132 (at 10 μmol/L) or BTZ (at 0.01-10 μmol/L), when compared with those stimulated by 0.5 μmol/L of MG132. This suggests that both 95-kDa and 85-kDa proteins may be also modified by ubiquitination or others.

### Putative ubiquitination of Nrf1α facilitates its proteolytic processing to yield a cleaved isoform of ^~^85-kDa

A previous report suggested that ubiquitination of Nrf1 occurs prior to, and is essential for, its proteolytic processing to yield several cleaved polypeptides [37]. However, our recent work has shown that putative ubiquitination of Nrf1α within its N-terminal domain (NTD) and adjacent acidic domain-1 (AD1) is not a necessary prerequisite for selective proteolytic processing of the protein to remover several N-terminal polypeptides during its maturation into an active CNC-bZIP transcription factor (data shown for another submission). To address this issue, all six lysines (as potential ubiquitin acceptors, Figure S7A) of Nrf1 within its N-terminal ubiquitin-like (UBL) domain (containing three lysines at positions 5, 6 and 70) and AD1 (containing other three lysines at positions 169, 199 and 205) were here mutated into another basic arginines. As anticipated, the resulting mutant Nrf1^6×K/R^ significantly diminished, but not completely abolished, production of the N-terminally-truncated protein-D (85-kDa) isoform, even in the presence of MG132 (at 5 μmol/L) (Figure S7B, *cf lanes 1-4 with 9-12*). By contrast, two major longer isoforms of protein-A (120-kDa) and -B (105-kDa) were unaffected by the mutant Nrf1^6×K/R^, but *in vitro* deglycosylation digestion by Endo H enabled both to convert into another two isoforms of protein-B (105-kDa) and protein-B^−^ (95-kDa), respectively (Figure S7B, *lanes 9-12*). Further examinations of single or double K/R-point mutants (Figure S7C) revealed that production of the processed 85-kDa protein-D of Nrf1 was also obviously prevented, as variable abundances (and stability) of its longer 120-kDa protein-A and/or 105-kDa protein-B were concomitantly, to more or less extents, decreased by Nrf1^K5/6R^, Nrf1^K70R^, Nrf1^K199/205R^, but not Nrf1^K169R^ or Nrf1^K199R^. These suggest that putative ubiquitination of Nrf1 through its N-terminal UBL-adjoining region appears to be involved in its proteolytic processing to yield the 85-kDa protein-D. This is further supported by another finding that yield of the processed 85-kDa protein-D was almost completely blocked by Nrf1^K56/70R^ (a mutant of all three lysines into arginines within putative UBL module), but not abolished by Nrf1^K169/199/205R^ (another mutant of all three lysines into arginines within the Neh2L region of AD1) (Figure S7E, *lanes 4 vs 12*). Intriguingly, after the recovery of single or double lysine-points from the Nrf1^6×K/R^ mutant was allowed for potential ubiquitination of Nrf1 at indicated lysines, the resultant yield (and stability) of the 85-kDa protein-D was still significantly diminished (Figure S7D). Similar results were obtained from other K/R-combined mutants (Figure S7E). This was accompanied by differentially altered abundances of the protein-A, -B and -B^−^ isoforms; they could be ubiquitinated even in some K/R mutant contexts.

### Nrf1α regulates key genes that are involved in its post-translational modification and proteolytic processing

After translation of the full-length Nrf1α protein, it is topologically integrated within and around the ER. Portion of its transaction domains (TADs) is partitioned in the lumen and glycosylated by oligosaccharyltransferase (as a enzymatic complex of distinct subunits such as those encoded by *OST4*, *STT3A* and *STT3B*). Subsequently, if required, the ER luminal-resident domain of Nrf1α is repositioned by p97-driven retrotranslocation machinery into extra-luminal side of membranes, where it is allowed for deglycosylation by peptide:N-glycosidase (encoded by *NGLY1*), ubiquitination by the ER membrane-bound ubiquitin ligase Hrd1 (also called SYVN1) [11, 36], and putative proteolytic processing by aspartic proteases DDI-1/2 [58, 59]. Thus, we examined which key genes required for post-translational modification of Nrf1α and its proteolytic processing are regulated by this CNC-bZIP transcription factor *per se via* a feedback loop. The results showed differential induction of some genes in different responses of between *Nrf1^+/+^* and *Nrf1α*^−/−^cells to stimulation by a lower (1 μmol/L) or higher (10 μmol/L) dose of MG132 (Figures 7 & S8). Interestingly, knockout of *Nrf1α*^−/−^ resulted in obvious decreases in both basal and stimulated expression of p97 at mRNA and protein levels (Figures 7F & S8B). However, significant increases in stimulated (but not basal) expression of Hrd1 and DDI-1 (Figures 7E,G and S8A) were found following treatment of *Nrf1α*^−/−^ cell with 1 μmol/L of MG132. Collectively, these results, together with other previously reported data [37, 69], demonstrate that there exist coupled positive and negative feedback circuits between Nrf1α and its target regulators (p97, Hrd1 and DDI), which are associated with the ER.

**Figure 7.**
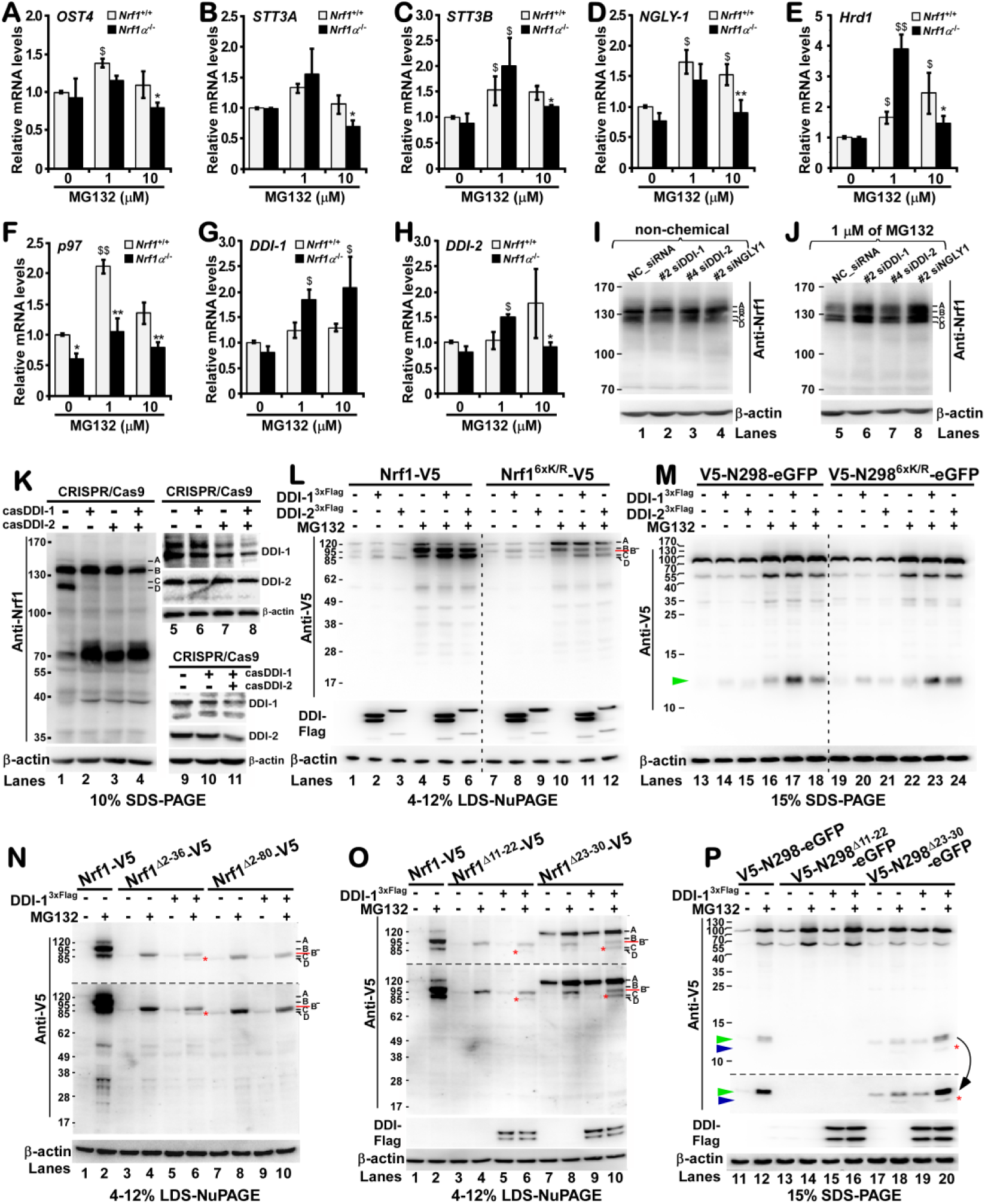
The proteolytic processing of Nrf1α by DDI-1/2 to yield some cleaved isoforms from 85-kDa to 12.5-kDa. **(A to H)** Two lines of HepG2 cells with distinct genotypes of wild-type *Nrf1^+/+^* and *Nf1α*^−/−^-specific knockout were treated with different doses of MG132 for 16 h before being lysates. Subsequently, both basal and MG132-stimulated mRNA expression of eight genes including *OST4*, *STT3A*, *STT3B*, *NGLY-1*, *Hrd1*, *p97*, *DDI-1* and *DDI-2* was determined by real-time qPCR. **(I,J)** Several siRNA sequences that had been selected by real-time qPCR in order to knock down expression of *DDI-1* (#2), *DDI-2* (#4) and *NGLY-1* (#2) (also see Figure S8 and Table S1), were transfected into HepG2 cells for 7 h, and allowed for a 16 h recovery culture. Then the cells were treated for additional 16 h with MG132 (1 μmol/L) or non-chemicals, and then examined by Western blotting with Zhang’s Nrf1 antibody. **(K)** HEK293 cells were subjected to disruption of *DDI-1* alone or plus *DDI-2* by both genomic editing by CRISPR/Cas9-mediated biotechnology. The resulting cell lines were identified by target gene sequencing and Western blotting with distinct antibodies against DDI-1, DDI-2 or Nrf1. **(L to P)** COS-1 cells were co-transfected for 7 h with each of a series of expression constructs for Nrf1, its mutants and its fusion proteins indicated with distinct tags, together with either DDI-1^3×Flag^ or DDI-1^3×Flag^, and then allowed for a 16 h recovery culture before being treated with MG132 (5 μmol/L) for 4 h prior to being lysates. Subsequently, these samples were subjected to protein separation by gradient NuPAGE gels containing 4-12% polyacrylamide (L,N,O) or general SDS-PAGE gels with upper and lower two-layers consisting of 8% and 12% polyacrylamide (M,P), followed by visualization by immunoblotting with antibodies against V5, Flag, or β-actin.

### The proteolytic processing of Nrf1α-derived proteins by DDI-1/2 proteases in close proximity of the ER

Knockdown of *DDI-1* or *NGLY1* by specific siRNAs (Figure S8, D to F) resulted in significant decreases in basal, but not MG132 (1 μmol/L)-stimulated, abundances of the processed Nrf1 protein-C/D isoforms (Figure 7I & J). This indicates that the putative MG132-triggered proteases, besides DDI-1, also contribute to proteolytic processing of Nrf1α to yield cleaved protein-C/D isoforms. Although knockdown of siDDI-2 at its mRNA levels appeared to have no effect on the processing of endogenous Nrf1 (Figure 7I), production of the cleaved Nrf1 protein-C/D isoforms was almost completely abolished by deficiency of DDI-2 alone or plus DDI-1 (i.e. both genes were partially deleted by CRISPR/ Cas9-mediated genomic editing, Figure 7K). Further, forced over-expression of DDI-2^3×Flag^, like DDI-1^3×Flag^, also led to marked increases in the abundance of cleaved ^~^85-kDa protein-D (C-terminally tagged by V5, Figure 7L). This cleaved isoform was generated from processing of ectopic Nrf1 or its mutant Nrf1^6×K/R^, albeit the mutant-derived isoforms of protein-B/B^−^ (105/95-kDa) and 85-kDa protein-D were reduced, when compared to equivalents from wild-type Nrf1 (Figure 7L, *lanes 10-12 vs 4-6*). Such being the case, it is thus postulated that a certain N-terminal polypeptide should be removed from Nrf1 or its mutant Nrf1^6×K/R^. As anticipated, an N-terminally V5-tagged ^~^12.5-kDa polypeptide was visualized (Figure 7M) by Western blotting of V5-N298-eGFP and mutant V5-N298^6×K/R^-eGFP (i.e. N298 indicates the N-terminal 298-aa of Nrf1, that is attached to the N-terminal end of eGFP). The yield of this ^~^12.5-kDa N-terminal polypeptide was significantly increased in the presence of DDI-1 or DDI-2 (Figure 7M, *lanes 16-18*, *and 22-24*).

In order to examine whether proteolytic processing of Nrf1α by DDI-1/2 proteases occurs in close proximity of the ER, HepG2 cells were co-transfected with expression constructs for DDI-1^3×Flag^ together with Nrf1^Δ2-36^-V5 (lacking the ER-targeting NHB1 signal peptide) or Nrf1^Δ2-80^-V5 (lacking both NHB1 and its adjacent CRAC peptide associated with ER membranes, but retaining NHB2). As shown in Figure 7N, co-expression of the Nrf1^Δ2-80^-V5 mutant with DDI-1^3×Flag^ only caused it to give rise to a single protein-D, which appeared to be exactly the faintly cleaved product of ^~^85-kDa from proteolytic processing of the major ^~^95-kDa Nrf1^Δ2-36^-V5 protein-B^−^ mediated by DDI-1^3×Flag^ (Figure 7N, *cf. lanes 6-10*), implying that a possible cleavage site exists within NHB2-adjoining peptides. Furthermore, a similar cleaved polypeptide of ^~^85-kDa was also generated from the proteolytic processing of the major ^~^95-kDa Nrf1^Δ11-22^-V5 protein (lacking the hydrophobic membrane-spanning h-region of the ER-targeting signal peptide) by co-expressed DDI-1^3×Flag^ (Figure 7O, *lane 6*). By contrast, an increased abundance of the cleaved ^~^90/85-kDa isoform arising from the proteolytic processing of the Nrf1^Δ23-36^-V5 mutant (lacking a luminal c-region of the ER-targeting signal peptide, which dictates its transmembrane-spanning orientation) by DDI-1^3×Flag^ was observed, even in the presence of MG132 (at 5 μmol/L) (Figure 7O, *lanes 10 vs 8*). The product increase was accompanied by a decrease of the longer ^~^95-kDa deglycoprotein of Nrf1^Δ23-36^-V5 but not its 120-kDa glycoprotein (*lanes 10 vs 8*). These demonstrate that the yield of the cleaved Nrf1 isoform is determined by orientation of its intact full-length protein topology integrated within the ER membranes.

It is interestingly to be found that an increased production of the N-terminal ^~^12.5-kDa polypeptide arising from the V5-N298^Δ23-36^-eGFP mutant was accompanied by a decreased abundance of another longer processed ^~^55-kDa intermediate form, but not of its full-length ^~^80-kDa protein (Figure 6P, *lanes 17-20*). By comparison with equivalents arising from wild-type V5-N298-eGFP, the processed ^~^55-kDa intermediate product was reduced by the mutant V5-N298^Δ23-36^-eGFP, but enhanced by another mutant V5-N298^Δ11-22^-eGFP (*cf. lanes 12 with 14*, *18*). However, no changes in abundance of the processed ^~^55-kDa isoform were observed upon co-expression of V5-N298^Δ11-22^-eGFP with DDI-1^3×Flag^, in the meantime while none of the N-terminal ^~^12.5-kDa polypeptide was also detected in this case (Figure 6P, *lanes 13-16*). These data indicate that generation of the N-terminal ^~^12.5-kDa polypeptide of Nrf1 from its fusion protein V5-N298-eGFP is dominantly dictated by its ER-anchored h-region (aa 11-22) within the NHB1 signal sequence. The output of the N-terminal cleaved ^~^12.5-kDa product is principally monitored by dynamic repositioning of signal peptide-associated sequence (aa 23-50), in cooperation with its adjacent membrane-tethered CRAC peptide (aa 55-80), which determines the proper topological orientation of Nrf1 within and around ER membranes, such as to control repartitioning of putative cleavage site-containing NHB2 peptide (aa 81-106) out of the organelle. However, it should be noted that deletion of the essential ER-anchored h-region conferred the resulting Nrf1^Δ11-22^-V5 mutant to substantially diminish, though not abolish, the yield of the minor C-terminal ^~^85-kDa polypeptide (Figure 7O, *lane 6*), but none of the cleaved N-terminal ^~^12.5-kDa polypeptide arises from V5-N298^Δ11-22^-eGFP (Figure 7P, *lanes 13-16*). The nuance might be attributable to a possible difference in topological locations of putative cleavage site-containing NHB2 along with the membrane-tethered CRAC peptide in distinct mutant contexts, albeit the detailed mechanism remains to be elucidated.

All together, these results demonstrate that the proteolytic processing of the ^~^95-kDa Nrf1 deglycoprotein is mediated by DDI-1 (and/or other MG132-triggered proteases) to remove its N-terminal ^~^12.5-kDa polypeptide, such as to yield an N-terminally-truncated ^~^85-kDa CNC-bZIP transcription factor. Such proteolytic processing of Nrf1 is also proposed to occur in close proximity to the ER. This notion is evidenced by the fact that upon deletion of the ER-anchored sequence and adjacent membrane-tether CRAC peptide (aa 2-80) to yield Nrf1^Δ2-80^-V5, not a cleaved isoform is detectable from this mutant co-expressed with DDI-1^3×Flag^, albeit it retains the putative DDI cleavage site (Figure 7N, *lanes 7-10*). In order to confirm which forms of Nrf1 are glycoprotein, deglycoprotein, and proteolytically processed protein, here we further examined COS-1 cells that had been transfected with an expression construct for V5-Nrf1-eGFP (in which the full-length Nrf1 is sandwiched by fusion with the N-terminal V5 ectope and the C-terminal eGFP). The results demonstrated that the p97 inhibitor NMS-873 caused a gradual accumulation of the fusion glycoprotein of V5-Nrf1-eGFP, but digestion by Endo H enabled this glycoprotein to disappear and be replaced by another deglycosylated protein (Figure S9). Both Nrf1-sandwiched glycoprotein and deglycoprotein were visualized by immunoblotting with antibodies against Nrf1 and its N-terminal V5 tag. However, the putative ^~^85-kDa Nrf1-fused protein with ^~^28.5-kDa eGFP, as well as additional N-terminally-truncated polypeptides, were detected by their cross-reactivity with anti-Nrf1 only, but not anti-V5, antibodies (Figure S9). Overall, these findings demonstrate convincingly that distinct lengths of the N-terminal portions (and adjacent regions) of intact Nrf1α is proteolytically truncated during its post-synthetic processing into a mature isoform of the active CNC-bZIP factor.

### Distinct processing of Nrf1α under distinct stresses induced by different concentrations of proteasomal inhibitor

Since Nrf1α is folded with dynamic membrane-topologies within the ER, it is reasonable to assume that topological folding of the CNC-bZIP protein and its post-translational processing should be, to greater or lesser extents, effected by distinct stresses induced by different concentrations of proteasomal inhibitors (e.g. MG132), particularly leading to an impairment of ERAD-dependent pathway. As anticipated, treatment of HepG2 cells with a lower (1 μmol/L) or higher (10 μmol/L) dose of MG132 conferred Nrf1 to be processed into distinct proteoform patterns (Figure 8A). This was accompanied by significant differences in accumulation of Nrf2 and its target antioxidant gene such as HO1, as well as an ER stress marker BIP. Further examinations revealed three classic ER stress-triggered signalling pathways, such as phosphorylation of elF2α (acting as a direct substrate of PERK), IRE1 (enabling XBP-1 mRNA to be spliced), and selective proteolytic processing of ATF6α, were differentially activated by different doses of MG132 (Figure 8B). The pulse-chase experiments unraveled distinct time- and/or dose-dependent effects of MG132 on activation of key signalling molecules within distinct ER stress-triggered UPR pathways (i.e. PERK-elF2α, XBP-1s and ATF6α-N, in Figure 8C). Overall, it is important to note that a lower dose (1 μmol/L) of MG132 only gave rise to a dominant stimulation signal to activate the PERK-elF2a pathway (Figure 8, C4 to C7), with a weak signal of XBP-1s but not of ATF6α-N. By contrast, striking signals of XBP-1s and ATF6α-N appeared to be activated by a higher dose (10 μmol/L) of MG132 (Figure 8, C8 & C9).

**Figure 8.**
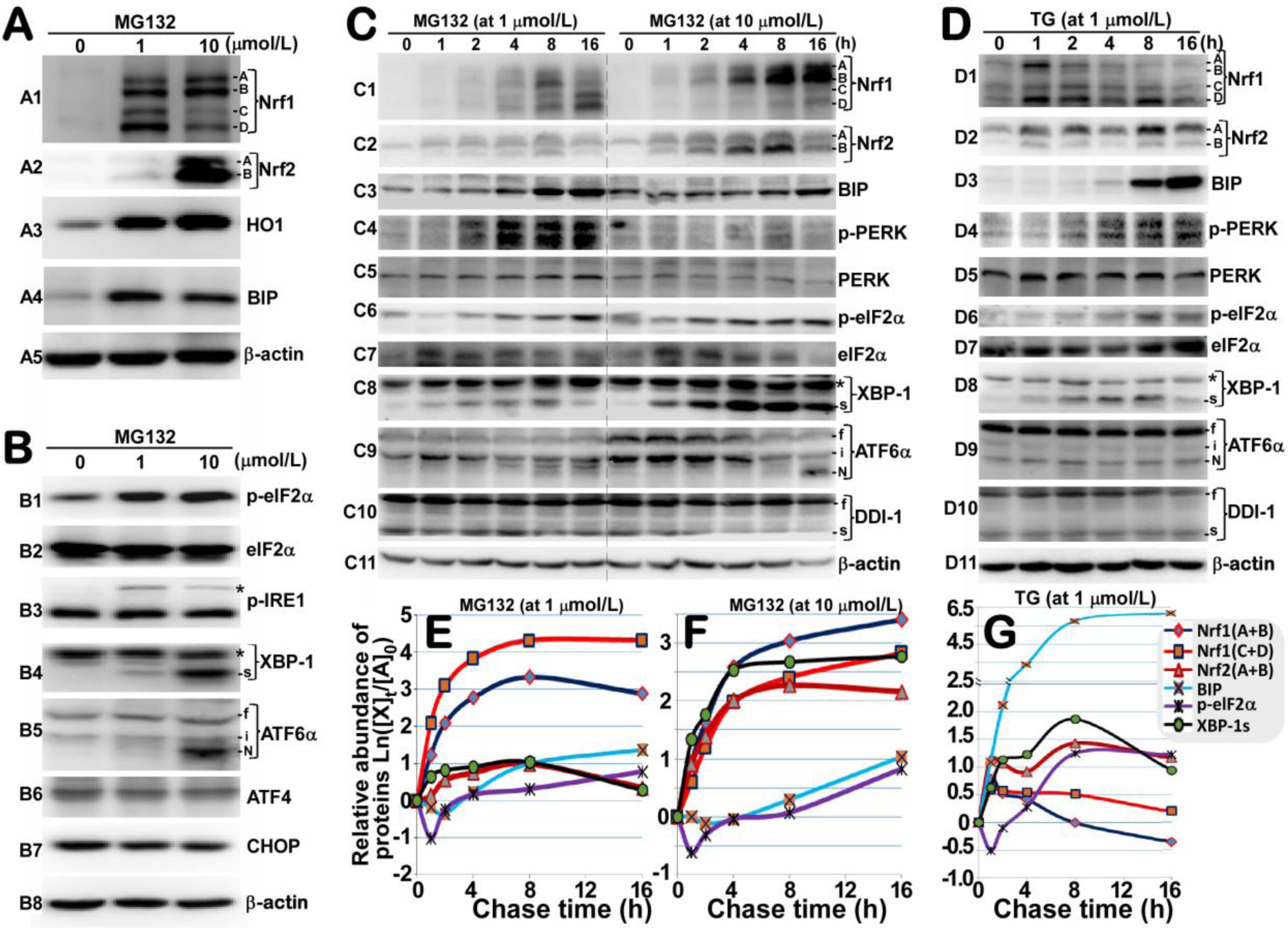
The post-synthetic processing of Nrf1α under grading ER stress induced by different doses of proteasomal inhibitors. **(A,B)** HepG2 cells were treated with a lower (1 μmol/L) or higher (10 μmol/L) dose of MG132 for 16 h, and then examined by Western blot with distinct antibodies against Nrf1, Nrf2 and other signalling molecules involved in the response to ER (and oxidative) stress. **(C to G)** Different concentrations of MG132 (1 μmol/L or 10 μmol/L) and TG (1 μmol/L) were administrated to treat HepG2 cells for distinct indicated times before being disrupted in denature buffer. The total lysates were resolved by SDS-PAGE gels and visualized by immunoblotting with distinct antibodies against Nrf1, Nrf2 or other indicated proteins (C,D). Subsequently, both basal and stimulated expression levels of major Nrf1 isoforms, as well as Nrf1 and other three signaling proteins (BIP, p-eIF2α and XBP-1s), were quantified and calculated as Ln(X) function values on the fold changes following treatment (E to G).

Stoichiometric analysis of distinct Nrf1 isoforms along with Nrf2 and UPR signaling molecules (such as BIP, p-elF2α and XBP-1s) illustrated there existed a possible correlation of different processing of Nrf1 under distinctive ER conditions of proteotoxic and oxidative stresses stimulated by different doses of MG132 (Figure 8, E & F). Such a similar relationship was, however, not observed following treatment of cells with TG (1 μmol/L, Figure 8D3); the ER stressor led to an incremental abundance of BIP (acting as a key chaperone sensor to ER stress and an effector of the resulting UPR), albeit this was accompanied by induction of other signaling molecules in a time-dependent manner (Figure 8, D & G). Interestingly, further comparison of the data obtained from stimulation of cells by MG132 or TG unveiled that the selective post-translational processing of the ER-located Nrf1 could also be affected by others (i.e. oxidative) besides ER stress. Together with the above results (as shown in Figure 3) and a previous report by another group [70], these findings demonstrate that distinct extents of oxidative stress and antioxidant response signaling are significantly induced by different concentrations of MG132, particularly at a higher dose of the proteasomal inhibitor, resulting in marked increases in the expression of Nrf2-target antioxidant genes (Figure 9). In addition, it is intriguing to note that the aspartic protease DDI-1 *per se* also appeared to be processed under modest ER stress induced by 1 μmol/L of TG or MG132 (Figures 8C10, 8D10, S8C). The putative processing of DDI-1 was also inhibited by a higher dose (10 μmol/L) of MG132 (as it stimulates a severe proteotoxic and oxidative stress with Nrf1 aggregation [55, 70, 71]). However, this warrants further studies to determine the mechanism(s) whereby distinct dose-dependent effects of proteasomal inhibitor on putative DDI-1 processing might contribute to limitation of proteolytic processing of Nrf1 under distinctive stress conditions.

**Figure 9.**
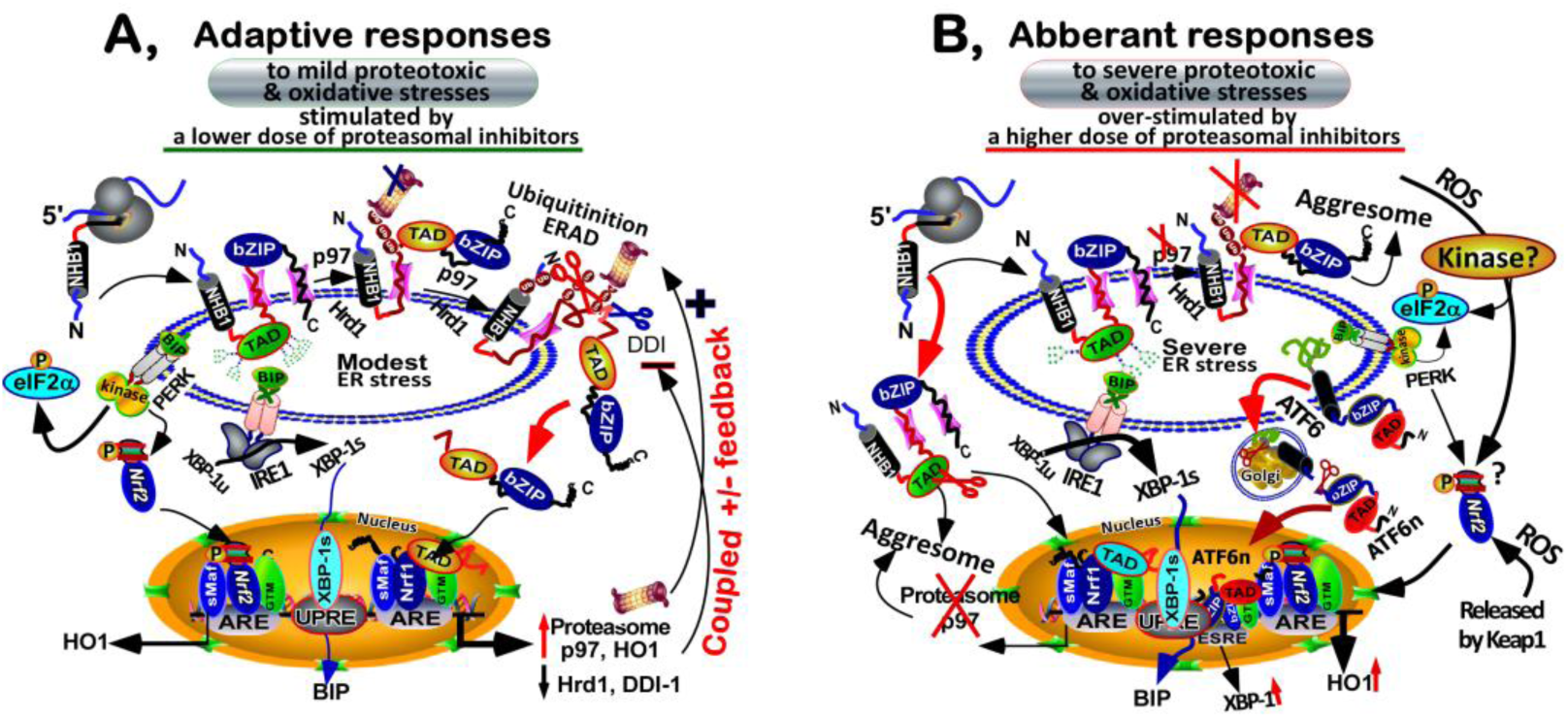
Coupled positive and negative feedback circuits existing between Nrf1 and its downstream target genes. Two proposed models are provided for a better understanding of molecular and cellular mechanisms that controls multistage post-translational processing of endogenous Nrf1 to yield distinct isoforms, which are involved within coupled positive and negative feedback circuits existing between the Nrf1 CNC-bZIP factor (alone or in cooperation with another homologous Nrf2) and its downstream target genes (some products of which also function just as its upstream regulators). Furthermore, discrepant topovectorial processing of Nrf1 to yield distinct isoforms within disparate subcellular compartments is affected and also involved in diverse ER-stress responses to a lower (H) or higher (I) concentration of proteasomal inhibitors, albeit detailed mechanisms remain to be determined.

## DISCUSSION

For undeniable reasons that largely involve difficulty of experimentation with unavailable models, the overwhelming majority of researchers in the field have been entirely fixated on Nrf2 (to the tune of thousands of publications per year), whilst Nrf1 appears to be comparatively ignored (with only a small number of publications [9]. As such, Nrf1 is identified to be actually indispensable for regulating critical cellular homeostatic and developmental processes, as well as adaptive cytoprotective responses. The main discoveries made have predicted a promising tendency for Nrf1 to become increasingly significant, which will attract much attentions from distinct related fields, particularly with the demonstration that it is a dominant regulator of transcriptional expression of 26S proteasomes and p97 through a positive feedback circuit, in intracellular response to challenges with proteasomal inhibitors as chemotherapeutic drugs [11, 35, 37, 46, 71]. Importantly, most of the intracellular Nrf1 (rather than Nrf2), with several domains being more conserved with the ancestral homologues CNC and Skn-1 [39, 72–74], is topologically integrated within the ER, before being sorted out of membranes and then translocated into the nucleus. This raises a question of how Nrf1 is regulated in distinct topovectorial processes from the ER to the nucleus. Transcriptional expression of Nrf1-target genes was finely tuned by distinct isoforms of the endogenous CNC-bZIP factor, such as to exert essential functions for maintaining cellular homeostasis and organ integrity, in the cytoprotective responses against a variety of cellular stresses from changing environments.

### Establishment of a generally acceptable criterion to identify endogenous Nrf1α/TCF11-derived isoforms

In this study, a large bulk of laborious work has been focused on endogenous Nrf1α/TCF11 and its post-translational processing, aiming to establish a general criterion acceptable for identification of multiple isoforms between 140-kDa and 100-kDa (Table 1). The objective of building such a generally acceptable criterion is for an attempt to terminate the chaotic state of the literature on distinct Nrf1α/TCF11 isoforms, which were defined confusedly by discrepant molecular weights of their endogenous proteins as inconsistently reported by those groups [27, 29, 35, 37, 55, 57, 61, 70]. For example, immensely disparate sizes of Nrf1α-derived proteoforms had been shown in three publications by Sha and Goldberg [37, 55, 71] (as the cropped images, Figure S1C to E). Of note, neither available positive standards of known Nrf1α/TCF11 *per se* nor Nrf1α/TCF11-specific knockout cells as convincing negative controls were shown together. Although the authors compared Nrf1 to distinct molecular weight standards and other reference proteins such as p97, p110, p85 and p105 [55], it is still so hard to distinguish these reference proteins from Nrf1α in diverse electrophoretic behaviors. This is owing to the fact that human Nrf1α of 742-aa, though it predicted to be an acidic protein (pI 4.57, DE/KR=113:65) with a mass of 81.5-kDa only, is *de facto* post-translationally modified with various mobility such as to locate between 140-kDa and 100-kDa, which are determined with its internal identifiable positive and negative controls required critically for different experimental settings (as referenced herein).

**Table 1.** Distinct isoforms of Nrf1 identified within different electrophoretic locations

| SDS-PAGE (8%+12%) | LDS-NuPAGE (4-12%) | Salient features |
| --- | --- | --- |
| Protein-A, ~140/145-kDa | ~120-kDa glycoprotein | N-glycosylation of Nrf1, that is inhibited by TU |
| Protein-B, ~130/135-kDa | ~95-kDa full deglycoprotein (B<sup>-</sup>);<br>~105-kDa partial deglycoprotein (B);<br>~95-kDa non-glycoprotein (TU-led) | Full deglycosylation of Nrf1 glycoprotein-A;<br>Partial deglycoprotein-B is transient unstable;<br>TU-led non-glycosylated nascent polypeptide |
| Protein-C & D, ~125/120-kDa | ~90/85-kDa processed protein | Proteolytical processing of Nrf1 to remove its<br>N-terminal ~12.5-kDa part of deglycoprotein-B/B<sup>-</sup><br>such as to yield a mature C-terminal ~85-kDa form |
| Protein-E & F, ~115/110-kDa | ~75-kDa, an N-terminally truncated<br>polypeptide examined somewhere | Proteolytical removal of its N-terminal domain<br>(NTD) plus an adjacent portion of AD1 from Nrf1;<br>Or may also be translated by an Nrf1<sup>ΔN</sup> variant |
| Short Nrf1β, ~65/70-kDa | ~55-kDa, a truncated isoform lacks<br>both NTD and AD1 regions of Nrf1 | Proteolytical processing to remove NTD & AD1;<br>Or may also be translated by an Nrf1β variant |

**Note:** Distinct Nrf1α -derived isoforms are resolved by two different electrophoresis systems (with distinct sample-denatured and gel-running buffers) and then visualized by immunoblotting with distinct antibodies against different regions of Nrf1 and the V5 tagged. Amongst them, **three Nrf1-specific antibodies are here used** as follows: (i) H285 (anti-Nrf1 from Santa), which recognizes the central peptide (aa 191-475) of human Nrf1/TCF11 (with its catalog No. sc-13031, from Santa Cruz Biotechnology Inc, Figure S1A, right panel); (ii) D5B10 (anti-Nrf1 from CST), which is a monoclonal antibody against human Nrf1/TCF11 at its N-terminal G129-surrounding peptide that is not overlapped with the adjacent aa 191-475 peptide (with its catalog No. 8052S from Cell Signaling Technology Inc, Figure S1A, left panel); and (iii) Zhang's anti-Nrf1 antibodies against its short isoform Nrf1b (aa 292-741, raised in the Zhang's own laboratory). Overall, only Nrf1α -derived longer isoforms have cross-reactivity with antibodies H285 and D5B10, whereas all those CNC-bZIP-containing proteoforms are recognized by our own Nrf1 antibodies.

Both endogenous and ectopic Nrf1α/TCF11-derived proteins expressed in experimental cell lines were resolved by two different electrophoresis systems (i.e. SDS-PAGE and LDS-NuPAGE). As is generally accepted, the migration of protein bands on electrophoresis gels is influenced by many factors including polyacrylamide concentration, pH value, and buffer type. In fact, protein separation should be also affected by distinct detergent types, which is attributed to the fact that lithium is a stronger cation than sodium, such that LDS should be a more potent detergent than SDS and thus exert a potent effect on protein denaturation. Overall, these lead to distinctive separation by between SDS-PAGE and LDS-NuPAGE gels of distinct Nrf1α/TCF11-derived proteoforms as defined in Table 1.

Nrf1/TCF11-expressing cells are firstly harvested in a mild lysis buffer (0.5% SDS, 0.04 mol/L DTT, pH 7.5), followed by deglycosylation reactions with glycosidase. The lysates are denatured in 1× LDS-NuPAGE sample loading buffer (pH 8.4) before addition of antioxidant reagent, and then separated by LDS-NuPAGE Bis-Tris gels containing 4-12% polyacrylamide in the MES-SDS running buffer (pH 7.3) at 150 V for 90-120 min at room temperature. Subsequently, western blotting of distinct Nrf1/TCF11 isoforms (with no matter which antibodies against Nrf1 or V5 tagged were blotted herein) clearly showed its full-length glycoprotein of ^~^120-kDa, along with its deglycoprotein of ^~^95-kDa and a shorter N-terminally-truncated isoform of ^~^85-kDa. Notably, our evidence has been provided revealing that the proteolytic processing of the ^~^95-kDa Nrf1 deglycoprotein, rather than its ^~^120-kDa glycoprotein, by DDI-1/2 aspartic proteases gave rise to a mature processed 85-kDa isoform (which may also be represented by two close proteaforms resembling cleaved isoforms of Nrf1 protein-C and -D, but the protein-C should be transient unstable to be rapidly degraded and disappear on some occasions).

By contrast, total cell lysates saved in the loading buffer (62.5 mmol/L Tris-HCl, pH 6.8, 2% SDS, 10% Glycerol, 50 mmol/L DTT, 0.1% Bromphenol Blue) were subjected to protein separation by classic SDS-PAGE gels that were made of two layers containing 8% and 12% polyacrylamide respectively, in the routine running buffer (25 mmol/L Tris-HCl, pH 8.8, 192 mmol/L Glycine, 0.1% SDS). The SDS-PAGE gels were allowed for the first 30-min running at 50 V, and thereafter the voltage was changed into 110 V until the time of continuously running was extended to ^~^150 min. Subsequently, western blotting with three distinct antibodies against Nrf1 and an additional V5 antibody revealed that at least four isoforms (called Nrf1 protein-A, -B, -C, and -D herein) were present, on most occasions, in over ten different cell lines examined. Both the full-length glycoprotein-A and grading deglycoprotein-B/B^−^ of Nrf1 are closely located between ^~^140-kDa and ^~^130-kDa, whilst two distinct lengths of processed Nrf1 protein-C and -D isoforms are migrated closely to ^~^120-kDa. Some portions of these findings appear to be also supported by a few lines of the previous evidence reported by [11, 70, 75]. Additional two smaller isoforms of protein-E and -F between ^~^110-kDa and ^~^100-kDa (that are more than molecular mass of Nrf1β between ^~^65-kDa and 70-kDa estimated on SDS-PAGE gels, whilst Nrf1β was also migrated to ^~^55-kDa on LDS-NuPAGE gels) were, on some occasions, examined in some indicated cell lines.

Collectively, the discrepancy in protein separation by between LDS-NuPAGE and SDS-PAGE gels demonstrates that distinct lengths of multiple Nrf1 isoforms with different post-translational modifications also conferred on them with various mobility during two different electrophoresis systems. By comparison of SDS-PAGE and LDS-NuPAGE gels in the present study and our past work [38, 39, 42, 44, 46], a greater advantage of NuPAGE has led us to recommend it for the study to determine the Nrf1 glycosylation status. However, SDS-PAGE gels containing 15% polyacrylamide, but not 4-12% LDS-NuPAGE gels, can enable a very small N-terminal polypeptide of ^~^12.5-kDa (which is inferable to arise from the selective proteolytic processing of within the N-terminal portion of Nrf1) to be detectably resolved.

### Mechanisms by which endogenous Nrf1α/TCF11 is post-translationally processed to yield multiple isoforms

Translation of Nrf1α/TCF11 gives rise to a newly-synthesized non-glycosylated polypeptide of ^~^95-kDa estimated on the LDS-NuPAGE gels, which is also resolved by SDS-PAGE gels as a protein-B isoform located closely to ^~^130-kDa, in particular treatment with TU. The N-terminal signal anchor peptide (of which the essential portion is called NHB1, aa 11-30) of this nascent protein enables it to be translocated through the Sec61 complex and integrated within and around the ER [11, 42, 44]. The NHB1-associated TM1 peptide (aa 7-26) of Nrf1 enables it to adopt a proper topology with an orientation of N_cyto_/C_Ium_ (i.e. its N- and C-terminal ends face the cytoplamic and luminal sides, respectively) spanning across membranes. Once the TM1 is anchored within membrane, its connective N-terminal UBL region [comprises a membrane cholesterol-binding CRAC sequence (aa 55-80) and a luminal-anchored NHB2 (aa 81-106)] and adjacent transactivation domains (TADs, containing AD1, NST and AD2 regions) are transiently repartitioned to entire the ER lumen where Nrf1 will be modified, whereas its DNA-binding CNC-bZIP domains are retained on the cyto/nucleoplasmic side of membranes.

Within the ER lumen, Nrf1 is allowed for N-linked glycosylation through its NST domain so as to become an inactive glycoprotein (of ^~^120-kDa estimated on the LDS-NuPAGE gels, which is also designated as protein-A isoform migrated closely to ^~^140-kDa on SDS-PAGE gels). If required, some portions of the luminal-resident domains within Nrf1 will be dynamically repositioned and retrotranslocated into the extra-luminal side of membranes, whereupon the protein is allowed for deglycosylation digestion to yield an active transcription factor (of ^~^95-kDa identified as a full deglycosylated protein). In addition, certain grading deglycoproteins between ^~^100-kDa and ^~^105-kDa were, on some occasions, examined by LDS-NuPAGE gels; they were also resolved by SDS-PAGE gels as two close protein-B/B^−^ isoforms located to 130-kDa. Further evidence has also been provided showing that, when N-linked glycosylation of Nrf1/TCF11 is blocked by TU, an OST-specific inhibitor, abundances of either its endogenous or stably-expressed glycoproteins are diminished gradually to disappear as the time of treatment with the chemical is extended to 2 h. Such changes in Nrf1 glycoproteins is accompanied by gradual decreases in the abundances of its deglycoprotein and processed protein-C/D until their disappearance. The protein-C/D forms are postulated to arise from the proteolytic processing of the N-terminal domain (NTD) in Nrf1 deglycoprotein-B/B^−^ rather than it glycoprotein-A, resulting in a concomitant removal of its smaller N-terminal ^~^12.5-kDa polypeptide. Additional progressive proteolytic removal of the UBL-adjoining AD1 in Nrf1 is inferable to give rise to other shorter protein-E & F isoforms (Table 1).

All together with the previous evidence provided by us and other groups from immunocytochemical staining of distinct Nrf1 isoforms, subcellular fractionation, and live-cell imaging combined with membrane protease protection assays [11, 39, 43, 44, 70], these demonstrated that protein-A and protein-B of Nrf1 are predominantly recovered in the ER-enriched membrane fractions, whereas minor protein-B and major protein-C/D are determined as the nuclear chromatin-binding factors. The protein-B and -C/D isoforms are also differentially distributed to exist in the cytosolic and nuclear soluble fractions. In addition to soluble fractions, those insoluble ubiquitinated (and/or carbonylated) Nrf1α/TCF11 and its aggregated proteins are also examined, particularly following treatment of cells with a higher concentration of proteasomal inhibitors [55, 70, 71].

Overall, these findings and others have convincingly demonstrated that, after Nrf1α/TCF11 biosynthesis, it has undergone a series of distinct successive selective post-translational modifications (e.g. glycosylation, deglycosylation and ubiquitination) and selective progressive processing by DDI-1/2 and other proteases to yield multiple isoforms in discrete subcellular locations. The variation of such proteoforms within different subcellular compartments is dictated by topovectorial processes of Nrf1. This is based on the fact that the nascent polypeptide loading through the Sec61 translocon enters the ER lumen where Nrf1α is glycosylated, subsequently it is folded into a proper membrane-topology with its distinct domains being positioned on the luminal or cytoplasmic sides of ER membranes. Thereafter, the luminal-resident portions of Nrf1α are partially repositioned through p97-driven retrotranslocation into the extra-ER cyto/nucleoplasms. Thereafter Nrf1 is allowed for deglycosylation by peptide:glycosidases (encoded by *NGLY-1*), ubiquitination by Hrd1, and proteolytic processing mediated by DDI-1/2 and MG132-ignited proteasome [37, 58, 59, 71]. This enable maturation of Nrf1 to yield an active CNC-bZIP factor, along with other proteaforms with distinctive or opposing activities, insomuch as to finely tune transcriptional expression of target genes (e.g. p97, Hrd1, DDI-1, and proteasomes), some of which may also, in turn, monitor the topovectorial processing of Nrf1α from the ER to the nucleus.

### The processing of Nrf1α/TCF11 within the directional coupled feedback response circuits to regulate target genes that are also induced by different concentrations of proteasomal inhibitors

The evidence has been here presented demonstrating that distinct extents of ER-derived proteotoxic and oxidative stresses were stimulated by different concentrations of proteasomal inhibitor MG132 (at 1-10 μmol/L). Inhibition of proteasome-mediated ERAD pathway leads to accumulation and even aggregation of ubiquitinated and carbonylated proteins (e.g. ER-localized Nrf1α/TCF11)[55, 70]. The proteotoxic stress is also accompanied by secondary oxidative stress leading to activation of Nrf2-mediate signalling. Further, a mild extent of ER-derived stress is stimulated by a lower dose (at 1 μmol/L) of MG132, leading to induction of the PERK-elF2α signalling pathway to attenuate general translation of nascent polypeptides, such that their loading to the ER is feedbackly alleviated. It is thus surmised that If the loading of Nrf1α/TCF11 into the ER was attenuated, yield of its glycoprotein-A, derived deglycoprotein-B and processed protein-C/D should be reduced or abolished. However, expression of glycoprotein-A and deglycoprotein-B of Nrf1 is *de facto* not terminated, whereas abundance of the processed Nrf1 protein-C/D is significantly incremented, upon partial inhibition of proteasomes by 1 μmol/L of MG132. Further evidence obtained from this study and others [11, 35, 37, 69] convincingly demonstrates that MG132-stimulated accumulation of Nrf1α/TCF11 protein-C/D, as a mature active transcription factor, leads to up-regulation of p97 and proteasomal subunits, but down-regulation of Hrd1 and DDI-1. Collectively, these indicate that there exist coupled positive and negative feedback circuits between Nrf1 and its cognate target genes, some of which may also be, in turn, involved in the successive post-translational modification and processing of this CNC-bZIP factor, in the intracellular ‘bounce-back’ response to relatively lower doses of proteasomal inhibitors.

Despite attenuation of general translation by ER stress-induced PERK-elF2a signalling, biosynthesis of Nrf1 and its proteolytic processing appear to be unaffected, but even enhanced, following treatment with a lower dose of MG132. Thus, a unique translational expression pattern of distinct Nrf1 isoforms is predicted to be controlled by its upstream open reading frame (uORF, which was analyzed by bioinformatics to exist within the upstream region of the intact full-length transcripts [9]), as described in the case of ATF4, c-Myc or C/EBP [76–78]. Subsequently, the post-synthetic Nrf1 should be subjected to a series of post-translational modifications and proteolytic processing in close proximity to the ER. These events are monitored by those feedback regulators encoded by Nrf1-target genes (i.e. proteasomal subunits, p97, Hrd1 and DDI-1). Amongst them, p97-driven retrotranslocation determines dynamic repositioning of Nrf1 into the extra-ER cyto/nucleoplasmic side of membranes, allowing its glycoprotein-A to undergo consecutive multistage processes, including deglycosylation by glycosidases, ubiquitination by Hrd1, and proteolysis by DDI-1 and other proteases (i.e. MG132-triggered proteasomes).

The yield of processed protein-C/D is significantly diminished by knockdown of DDI-1-targeting siRNA, but the diminishment is completely abolished by 1 μmol/L of MG132, implying that the protein-C/D arises from proteolytic processing of Nrf1 by DDI-1 and/or other DDI-independent proteases (i.e. MG132-induced proteasomes). This notion is further supported by in-depth insights into putative proteasome-mediated processing of Nrf1 [71], demonstrating that either site-specific proteasomal inhibitors in a dose-dependent manner or CRISPR-mediated genetic inactivation of β1/PSMB6, β2/PSMB7 and/or β5/PSMB5 significantly suppress production of a soluble, active Nrf1 isoform (which is similar to the processed protein-C/D) and hence prevent the recovery of proteasome activity through induction of *de novo* proteasomes by its partial inhibition. Further studies by us and others [37, 55, 71] reveal that stimulated accumulation of the processed Nrf1 protein-C/D by a lower dose of MG132 is strikingly reduced, but not completely abolished, by sufficient inhibition of proteasomes by a higher dose (10 μmol/L) of the same chemical. Such a higher dose of proteasomal inhibitor stimulates severe ER-derived proteotoxic and oxidative stresses, leading to aggregation of Nrf1 protein-A/B (as are ubiquitinated and carbonylated [55, 70, 71]). The putative processing of Nrf1 may be thus prevented by complete inhibition of proteasomes by higher doses of MG132, albeit detailed mechanism(s) remains elusive. Moreover, two primary UPR signalling pathways mediated by IRE1-XBP1 and ATF6 are also activated by 10 μmol/L of MG132, but it is unknown whether they are involved in Nrf1 processing.

The proteolytic processing of ectopic Nrf1α by DDI-1/2 to yield a processed protein of ^~^85-kDa (estimated by NuPAGE) is not impaired, but conversely incremented, by sufficient inhibition of proteasomes by 5 μmol/L of MG132. Such MG132 treatment of Nrf1^Δ23-30^-expressing cells only caused the mutant (lacking the c-region of ER-anchored signal peptide that determines the proper topological orientation of Nrf1 within membranes) to yield a fainter processed isoform of ^~^85-kDa, but its production is markedly enhanced by DDI-1, which mediates the proteolytic processing of deglycoprotein and/or non-glycoprotein of ^~^95-kDa (estimated by NuPAGE). In addition, only a little of the ^~^85-kDa protein emerged following 5 μmol/L of MG132 treatment of cells, allowed for co-expression of DDI-1 with either Nrf1^Δ2-36^-V5 (lacking entire ER-targeting signal peptide) or Nrf1^Δ11-22^-V5 (lacking essential ER-anchored h-region), but not Nrf1^Δ2-80^-V5 (lacking the ER-targeting and cholesterol-binding CRAC sequences). Together, these findings demonstrate that the processed ^~^85-kDa Nrf1 protein arises from proteolytic processing of its ^~^95-kDa deglycoprotein precursor by DDI-1/2 in close proximity to the ER, and also is further targeted to the ERAD-dependent proteolysis mediated by proteasomes.

### Concluding remarks

The objective of the present study is to establish a general criterion acceptable for identification of the endogenous Nrf1α/TCF11 and derivative isoforms, with distinct masses and half-lives determined in distinct experimental settings. Herein, we have elucidated molecular mechanisms that dictate successive multistate post-translational modifications (i.e. glycosylation by OST, deglycosylation by NGLY, and ubiquitination by Hrd1) of Nrf1 and its proteolytic processing to yield multiple proteoforms. Several lines of experimental evidence demonstrate that the nascent polypeptide (i.e. non-glycoprotein) of Nrf1α/TCF11 is transiently translocated into the ER, in which it is glycosylated to become an inactive glycoprotein-A, and also folded in a proper topology within and around membranes. Subsequently, dynamic repositioning of the luminal-resident domains in Nrf1 glycoprotein is driven by p97-fueled retrotranslocation into extra-ER cyto/nucleoplasms, allowing its glycoprotein-A to be digested by glycosidases into an active deglycoprotein. The progressive proteolytic processing of Nrf1 deglycoprotein-B/B^−^ by DDI-1/2 and other proteases (e.g. proteasomes) to give rise to a mature protein-D and other processed isoforms, which is accompanied by removal of its N-terminal ^~^12.5-kDa and other polypeptides. Interestingly, this study also unravels that coupled positive and negative feedback circuits exist between Nrf1 and its target genes (encoding its feedback regulators p97, Hrd1, DDI-1 and proteasomes). These key players are differentially and even oppositely involved in discrete cellular signalling responses to distinct extents of both ER-derived proteotoxic and oxidative stresses, induced by different doses of proteasomal inhibitors. Further evidence has been provided for a better understanding of mechanisms that control dual opposing effects of proteasomal inhibitors on differential expression of Nrf1, that is activated in the intracellular ‘bounce-back’ response to lower doses of proteasomal inhibitors, but is also suppressed by higher doses of the same chemicals.

## MATERIALS AND METHODS

### Chemicals and antibodies

All chemicals were of the highest quality commercially available. The protein synthesis inhibitor cycloheximide (CHX), the proteasome inhibitor MG132 and Calpain inhibitors (ALLN/CI, CII and CP) were purchased from Sigma-Aldrich (St Louis, MO, USA). Another two proteasome inhibitors bortezomib (BTZ) and epoxomicin (Epox) were obtained from ApexBio (USA). Tunicamycin (TU) and thapsigargin (TG) were from Sangon Biotech (Shanghai, China). NMS-873, a p97 specific inhibitor, was from Selleck (USA). Three antibodies against endogenous Nrf1 proteins were acquired from Cell signaling (D5B10, Cat# 8052S), Santa Cruz (H-285, Cat# sc-13031) and our lab (i.e. Zhang’s indicated in this study [44]), respectively. Moreover, other antibodies against the V5 ectope (Ivitrogen, Shanghai, China), the Flag (Beyotime Biotechnology, Shanghai, China), Ub (Cell Signaling Technology, Bedford, MA, USA), p97 (Abcame, Shanghai, China) or Hrd1 (ABclonal, USA) were employed, whilst β-actin and secondary antibodies were from ZSGB-BIO (Beijing, China).

### Cell culture and transfection

Several lines of HepG2, MHCC97L, HL7702, and HEK293 were originated from ATCC (Zhong Yuan Ltd., Beijing, China). RL-34 and COS-1 cell lines were maintained from Prof. John D. Hayes’ lab (University of Dundee, UK). MEFs with wild-type *Nrf1^+/+^* and its knockout mutant (*Nrf1*^−/−^) were provided as a gift by Dr. Akira Kobayashi (Doshisha University, Japan). SH-SY5Y cell line was friendly from by Prof. Zezhi Wu (Chongqing University, China). Both JB6 and GC-2 cell lines were kindly gifted by Prof. Julia Li Zhong (Chongqing University, China). The Rat tendon primary cells were generously gifted by Dr. Chuanchuan Lin (Southwest hospital, Chongqing, China). Nrf1α-specific knockout (Nrf1α^−/−^) HepG2 cell line was constructed in our lab by Dr. Yonggang Ren [48], whilst Nrf2-specific knockout (Nrf2^−/−^) in HepG2 cells, Nrf1α^−/−^ made in HL7702 cell line, HEK293-C (i.e. tetracycline-inducibly stable expression of human full-length Nrf1) and sh-Nrf1 (i.e. short-hairpin RNA to knock down Nrf1 in HepG2 cell lines were established in our lab, by using distinct biotechnologies, such as CRISP-Cas9, TALENs, Flp-In^TM^ T-REx^TM^-293 cell and lentivirus packaging systems (for detailed information unpublished) respectively. Most of these cell lines were allowed for growth in DMEM basic (1X) medium (GIBCO, Life technologies), with an exception of SH-SY5Y that was cultured in RPMI 1640 medium (GIBCO, Life technologies), which were supplemented with 10% (v/v) foetal bovine serum (FBS, Biological Industries, Israel), 100 units/ml of penicillin and streptomycin, in the 37°C incubator with 5% CO2. Equal numbers of experimental cells were seeded into each of 35-mm culture dishes, and then when the cells reached to 60%^~^80% confluence, they were transfected for 7 h with siRNAs and/or indicated expression plasmids, by using the Lipofectamine-3000 (Invitrogen).

### Establishment of DDI-1/2-specific knockout cell line

The DDI-1/2-specific knockout cells were established by the CRISPR/Cas9-mediated genome-editing system in HEK293. The sequences of single guide RNA (gRNA) targeting specifically for DDI-1 and DDI-2 were designed to be represented by 5’-CACCGTGTACTGCGTGCGGA-3’ and 5’-GGTCGACGCCGACTTCGAGC-3’, respectively, through the software CRISPR-direct (http://crispr.dbcls.jp/) and inserted into the pCAG-T7-cas9+gRNA-pgk-Puro-T2A-GFP vector (Viewsolid Biotech, Beijing, China). After transftection into HEK293 cells, the positive clones were selected by resistance to puromycin (2.5 mg/mL) and then confirmed by Western blotting, before being employed in some experiments.

### Luciferase reporter assay

Equal numbers (1.0×10^5^) of experimental cells were seeded into each well of the 12-well plates. After reaching 80% confluence, the cells were transfected with a Lipofectamine 3000 mixture with *P_SV40_GSTA2-6×ARE-Luc*, which contains six copies of the core ARE consensus sequence from rat GSTA2, or *3×PSMA4-ARE-Luc* that contains three copies of the core ARE from the promoter of *PSMA4* [46], along with a *Renilla* reporter as an internal control for transfection efficiency. The ARE-driven Luciferase activity was measured by using the dual-luciferase reporter assay system (E1910, Promega), at 30 h after transfection of cells that were allowed for additional 16-h treatment with different doses of proteasome inhibitors. The basal and stimulated ARE-driven reporter activity mediated by endogenous Nrf1/2 was calculated in its values against the background levels, and then the resultant data were further normalized as a fold change (mean ± S.D), relative to the basal activity of untreated group (at a given value of 1.0). The data presented here represent at least 3 independent experiments undertaken on separate occasions that were each performed in triplicate. Significant differences in the transcriptional activity were subjected to statistical analysis.

### Expression constructs and siRNAs

A series of the expression constructs for distinct mutants corresponding to different lysine (K) positions at 5, 6, 70, 169, 199, 205 into arginine (R) residues within NTD and AD1 of Nrf1 were made as described previously [40]. Further, both DDI-1 (Gene ID: 414301) and DDI-2 (Gene ID: 84301) were cloned from HEK293 cells and then inserted into the KpnI/BamHI site of the p3×Flag-CMV-14 vector. All oligonucleotide primers were synthesized by TSINGKE Biological Technology (Chengdu, China), and the fidelity of all cDNA products was confirmed by sequencing.

In order to knock down expression of the indicated genes, their small interfering RNA sequences targeting against the human DDI-1, DDI-2, NGLY1 (Gene ID: 55768), p97 (Gene ID: 7415) and Hrd1 (Gene ID: 84447) were synthesized by BIOTEND (Shanghai, China), as follows: siDDI-1 #1: 5’-CAUGAAUAUAGCGAUAGAA-3’, #2: 5’-GUCAUGGAUUCAGGA CGAA-3’, #3: 5’-CAAGUGACGAUGCUCUACA-3’; siDDI-2 #4: 5’-GUGCCCAGAUGACUAUCAU-3’, #5: 5’-GAGAUAUGUUGC UGGCCAA-3’, #6: 5’-CUCAUCUCCUGGAGAAAUA-3’; siNGLY1#1: 5’-GCCACUCCUUUAUGAAAUA-3’, #2: 5’-GCUCCACCUU UGUUACAAU-3’, #3: 5’-GCACUGGUUUAAGGAAGAA-3’; si-p97 #1: 5’-GCCAAUGUCAGAGAAAUCU-3’, #2: 5’-GGGCACA UGUGAUUGUUA-3’, #3: CCGUCGAGAUCACUUUGAA-3’; and si-Hrd1 (5’-UGGAGGAGGCAGCAGCAACAACUGU-3’). In addition, a scrambled siRNA sequence (5’-UUCUCCGAACGUGUCACG-3’), with no homology with any genes analyzed in Genbank, served as an internal negative control.

### Quantitative real-time PCR analysis

HepG2 cells that had been transfected with the indicated siRNAs, were subjected to isolation of total RNAs, according to the instructions of the RNAsimple Total RNA Kit (from Tiangen Biotech CO., LTD, Beijin, China). Subsequently, 500 ng of total RNAs was allowed for generation of the first strand of cDNAs (by the RevertAid First Strand Synthesis Kit from Thermo), which served as the template of quantitative PCR in the GoTaq^®^ qPCR Master Mix (promega). Then, each pairs of forward and reverse primers (Table S1) were added in the PCR, that was performed in the following conditions: inactivation at 95°C for 10 min followed by 40 cycles of 15s at 95°C and 30s at 59°C. At last, the melting curve was added to examine the amplification quality. In addition, expression of β-actin or RPL13A mRNAs was used as an internal control for normalization.

### *In vitro* deglycosylation reactions

The *in vitro* deglycosylation reactions of each sample with 500 units of Endo H (New England Biolabs #P0702L) were carried out as described previously [42] and then visualized by Western blotting.

### Western blotting

Experimental cells were transfected with distinct expression constructs for about 7 h, and then allowed for a 16-h recovery in DMEM containing High glucose. The cells were treated with chemicals for indicated times before being harvested at 4°C in a lysis buffer (0.5% SDS, 0.04 mol/L DTT, pH 7.5) containing the protease inhibitor EASYpacks (Roche, Germany). The samples were immediately denatured at 100°Cfor 10 min and sufficiently sonicated, prior to being allowed for reduction reaction with 3× loading buffer (187.5 mmol/L Tris-HCl, pH 6.8, 6% SDS, 30% Glycerol, 150 mmol/L DTT, 0.3% Bromphenol Blue) or 1× LDS-NuPAGE sample loading buffer (pH 8.4, with antioxidant reagent, both were from Ivitrogen, Shanghai, China), at 100°C for 5 min. Equal amounts of prepared protein extracts were subjected to separation by SDS-PAGE containing 10% or 15% polyacrylamide in the Tris-glycine SDS running buffer (pH 8.3) or LDS-NuPAGE gel containing 4% to 12% polyacrylamide with MES SDS running buffer (pH 7.3, from Ivitrogen, Shanghai, China). The resolved proteins were transferred onto the PVDF membranes, which were blocked by incubation with 5% non-fat milk at room temperature for 1 h before additional incubation with the primary antibodies for overnight at 4°C and cross-reaction for 2 h with the secondary antibodies. The immunoblots were developed with SuperSignal West Femto maximum sensitivity substrate (from Thermo) and then visualized by automatic imaging system (i.e. VersaDoCTM-4000MP from Bio-Rad) or MyECL imager (Thermo). On some occasions, the PVDF membranes blotted with antibodies were rinsed in stripping buffer for 30 min before being re-probed with an additional antibody, whilst β-actin served as an internal control to verify equal loading of protein into each of electrophoretic wells. The intensity of immunoblots was quantified by using the Quantity One software.

### Statistical analysis

Statistical significances of fold changes in the *GSTA2-6×ARE-Luc*, *3×PSMA4-ARE-Luc* reporter activity and also in the gene expression were determined using the Student’s t-test or Multiple Analysis of Variations (MANOVA). The data are shown as a fold change (mean ± S.D), each of which represents at least 3 independent experiments that were each performed triplicate.

## Supplemental Information

Supplemental Information includes nine figures and one table.

## Footnotes

**Author contributions:** Y.X. constructed most of the plasmids except those indicated, performed most of Western blotting of endogenous Nrf1, measured some lucifurase assays, collected the data and prepared drafts of most figures. Both M.W. and S.H. performed some of Western blotting work on ER stress and qPCR, and prepared drafts of some figures. L.Q., F.Y. and Z.Z. made the invaluable cell lines and repeated the qPCR, lucifurase assays and other experiments. S.Y. and J.P. provided critical suggestion and other valuable information to improve the work. Y.Z. designed this study, analyzed all the data, prepared all figures, wrote and revised the paper.

## Acknowledgments

The study was supported by the National Natural Science Foundation of China (key programs 91129703, 91429305 and project 31270879) awarded to Prof. Yiguo Zhang (University of Chongqing, China), and in part funded by Chongqing University postgraduates innovation project (No. CYB15024) awarded to Mr. Lu Qiu.

